# PermaPhos^Ser^: autonomous synthesis of functional, permanently phosphorylated proteins

**DOI:** 10.1101/2021.10.22.465468

**Authors:** Phillip Zhu, Rachel Franklin, Amber Vogel, Stanislau Stanisheuski, Patrick Reardon, Nikolai N. Sluchanko, Joseph S. Beckman, P. Andrew Karplus, Ryan A. Mehl, Richard B. Cooley

## Abstract

Installing stable, functional mimics of phosphorylated amino acids into proteins offers a powerful strategy to study protein regulation. Previously, a genetic code expansion (GCE) system was developed to translationally install non-hydrolyzable phosphoserine (nhpSer), with the γ-oxygen replaced with carbon, but it has seen limited usage. Here, we achieve a 40-fold improvement in this system by engineering into *Escherichia coli* a biosynthetic pathway that produces nhpSer from the central metabolite phosphoenolpyruvate. Using this “PermaPhos^Ser^” system – an autonomous 21-amino acid *E. coli* expression system for incorporating nhpSer into target proteins – we show that nhpSer faithfully mimics the effects of phosphoserine in three stringent test cases: promoting 14-3-3/client complexation, disrupting 14-3-3 dimers, and activating GSK3β phosphorylation of the SARS-CoV-2 nucleocapsid protein. This facile access to nhpSer containing proteins should allow nhpSer to replace Asp and Glu as the go-to pSer phosphomimetic for proteins produced in *E. coli*.

## INTRODUCTION

Reversible protein phosphorylation events underlie the exquisitely orchestrated regulation of many signaling pathways, with imbalances in these systems being key signatures of disease.^1, 2^ In humans, over 75% of proteins are reversibly phosphorylated at one or more sites, with nearly 80% of these being phosphoserine (pSer) modifications,^3^ on which we focus here. Two factors have greatly limited efforts to define the interactions and functions of specific phospho-proteins: (1) the challenge of making specific homogeneous, and quantitatively phosphorylated protein forms for *in vitro* study and (2) the challenge of the transient nature of phosphorylation in the cellular milieu due to phosphatase activity^4, 5^. The first challenge has now been well-addressed through genetic code expansion (GCE) systems that enable efficient, translational incorporation of pSer at genetically programmed codons into recombinant proteins produced in *E. coli*.^6–8^ However, the second challenge remains.

In addressing the challenge of the transient nature of pSer sites, non-hydrolyzable (i.e. stable) analogs or mimics of phosphorylated amino acids (Fig. 1A) have been indispensable for *in vitro* and in cell studies, for creating inhibitors of phosphoprotein-binders, kinases and phosphatases that can serve as biochemical tools and potential therapeutics^4, 5, 9–11^, and for making stable antigens for raising antibodies to recognize specific phosphoproteins^12^. The practical utility of such phosphomimetics hinges both on how well they faithfully recapitulate the effects of phosphorylation, and on the ease with which they can be quantitatively incorporated into proteins of interest.

**Figure 1.**
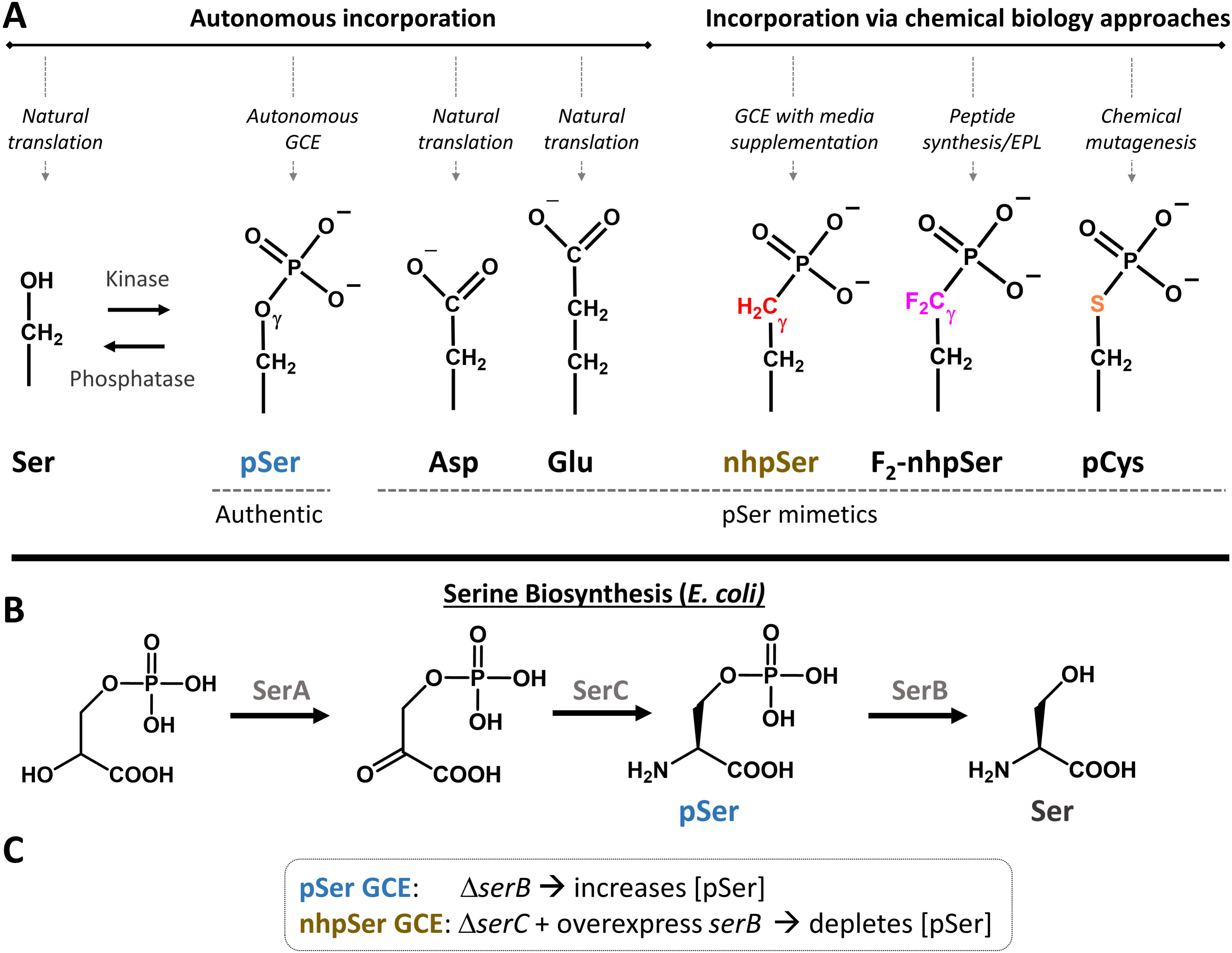
Mimics and biosynthesis of phosphoserine. (A) Phosphoserine and its mimics can be incorporated into proteins via a variety of techniques. Autonomous incorporation implies the free amino can be biosynthesized by the expression host and then incorporated into proteins during translation. Other mimics such as nhpSer, F_2_-nhpSer and pCys have previously only been installed using the indicated chemical biology approaches. (B) Phosphoserine is a biosynthetic intermediate of serine. (C) This pathway can be altered for GCE purposes to elevate (for pSer GCE) or eliminate (for nhpSer GCE) intracellular concentrations of the free phosphoserine amino acid.

The only widely employed pSer (and pThr) phosphomimetics are the natural amino acids Asp and Glu (Fig. 1A). Their main advantages are that they are trivial to encode into any protein for *in vitro* or *in vivo* studies, and they confer a negative charge at a roughly proper distance from the backbone that in some cases induces some functional changes similar to those caused by phosphorylation^13–15^. However, very well documented, even if generally underacknowledged, is that Asp and Glu are actually rather poor pSer mimics: their charge (-1 versus -2 for pSer at physiological pH), shape and hydrogen-bonding geometries are all markedly different than those of pSer,^16–18^ and indeed Ser to Asp/Glu mutations regularly fail to faithfully recapitulate the functional effects of phosphorylation (e.g.^19–22^).

Much better structural mimics than Asp and Glu are phosphono-methyl-alanine (referred to here as “non-hydrolyzable pSer” or nhpSer) and its difluorinated analog (F_2_-nhpSer) (Fig. 1A). In these isosteric mimics, the bridging γ-oxygen of pSer is replaced with a CH_2_ or CF_2_ group, respectively. Because the electronegative F-atoms of F_2_-nhpSer (i) can be H-bond acceptors like the pSer γ-oxygen and (ii) lower the phosphonate pKa2 from in the 7 to 8 range to near 5.5 so that its charge at physiological pH is ∼ -2 (like pSer), it is generally considered a better analog than nhpSer. In fact, reviews give the impression that nhpSer is not a useful mimic for these reasons^5, 23^, yet the effectiveness of both has been established in the few studies in which they were incorporated into synthetic peptides^12, 24–27^ and into proteins via semi-synthetic methods such as expressed protein ligation (EPL)^28–31^. For example, proteins Histone 2A^30^ and AANAT^28^ with nhpSer (but not Asp/Glu) installed at their N-termini by EPL were able to pull down 14-3-3, a pSer/pThr specific binding protein, from cellular extracts. Consistent with this, a nhpSer containing peptide bound 14-3-3 with comparable affinity to the equivalent pSer-peptide (0.86 vs 0.46 μM, respectively)^24^.

Despite their much more effective mimicry compared with Asp and Glu, both nhpSer and F_2_- nhpSer remain little-used due to the limited commercial availability of the compounds, the lack of an easy way to put them into proteins, and restrictions on which sites they can be installed by EPL. Just as GCE has provided a solution to efficiently producing authentic pSer proteins^6–8^, it offers an attractive strategy to incorporate nhpSer translationally and site-specifically into proteins at genetically programmed amber (TAG) stop codons. In fact, when in 2015 Chin and colleagues developed their high efficiency GCE approach to installing pSer into proteins, they showed that the same GCE components could incorporate exogenously added nhpSer in *E. coli*,^7^ and in 2018 they showed the system could be adapted for mammalian cells^32^. However, the *E. coli* system was neither optimized nor quantified for nhpSer incorporation^7^, and in the mammalian system less than 50% of the target protein contained nhpSer^32^. While we were finalizing this manuscript, the first use of this nhpSer *E. coli* GCE system outside the originating lab was reported, and it involved incorporating nhpSer (as well as pSer) into ubiquitin and NEDD8.^33^ These studies provided a further example of nhpSer being a more effective pSer mimic than Asp and Glu, and at the same time revealed shortcomings of the existing GCE system^7^ for making pure nhpSer proteins: high media concentrations of expensive nhpSer (8 mM) were needed to drive its incorporation, and even then the targeted nhpSer-protein products co-purified with both pSer-containing and non-phosphorylated protein. These limitations could explain why this GCE system has found limited use to generate nhpSer containing proteins.

Here, we hypothesize that the shortcomings of the above system for nhpSer incorporation into proteins is rooted not in a limitation of the GCE components, but simply in poor cellular uptake of the nhpSer amino acid from the media. Guided by the success of pSer and pThr GCE systems that use engineered *E. coli* to biosynthesize the required phospho-amino acid^6, 7, 34^, we engineered into *E. coli* a 6-step biosynthetic pathway that produces nhpSer from the central metabolite phosphoenolpyruvate. When coupled with the existing pSer GCE machinery, this system – which creates a self-sufficient, 21-amino acid *E. coli* GCE expression system we call “PermaPhos^Ser^” – efficiently generates milligram quantities of full-length, homogenous, permanently phosphorylated proteins containing nhpSer. Finally, to address the question of the efficacy of nhpSer as a pSer mimic, we use proteins produced by PermaPhos^Ser^ to measure the pKa2 values of nhpSer and pSer in a protein context, and to carry out direct comparisons of three nhpSer- and pSer-containing proteins and document that nhpSer faithfully recapitulates the functional effects of serine phosphorylation in multiple stringent systems where Asp/Glu mimetics could not.

## RESULTS

### Evaluation of the existing nhpSer GCE system

Chin and colleagues reported that by eliminating intracellular pSer (using an *E*. coli Δ*serC* mutant and over-expressing *serB* to hydrolyze pSer that might get into the cell, Fig. 1B and 1C) and supplementing the media with nhpSer they were able to install nhpSer at programmed amber (TAG) stop codons using the same amino-acyl tRNA synthetase (aaRS)/tRNA pair and EF-Tu (EF-Sep) that allowed pSer incorporation.^7^ Our first goal was to quantify the efficiency of that system by using it to incorporate either one or two nhpSer residues into a superfolder green fluorescent protein (sfGFP) reporter protein. Using their conditions of 2 mM nhpSer in the media culture, fluorescence values obtained were slightly above those of cultures lacking nhpSer for single TAG codon suppression (corresponding to ∼1-2 mg per liter culture), but the double TAG codon suppression cultures produced indistinguishable quantities whether nhpSer was added or not (Supplemental Fig. 1). Whatever the origin of the low background fluorescence signal observed in cultures lacking nhpSer (perhaps due to small quantities of full-length sfGFP produced via near-cognate suppression), Phos-tag gel electrophoresis^35^ of purified protein produced from the single TAG site suppression was consistent with mostly nhpSer-containing protein with some contamination with pSer-containing protein (Supplemental Fig. 1)^32^. This confirms that the published GCE system is indeed functional for nhpSer incorporation, but it gives low yields and some contamination with pSer-containing protein that would be difficult to purify away. We reasoned this low efficiency was due to low bioavailability of the nhpSer amino acid. GCE systems for pSer and pThr incorporation overcame this issue by leveraging biosynthetic pathways that produce high intracellular phospho-amino acid concentrations,^6–8, 34^ and so we sought to do the same for nhpSer.

### Developing an efficient nhpSer-protein expression system

#### A putative nhpSer biosynthetic pathway

A putative biosynthetic pathway for nhpSer was fortuitously discovered in 2010, when Zhao and colleagues probed the 10-step pathway by which *Streptomycess rubellomurinus* converts phosphoenolpyruvate into the anti-malarial natural product FR-900098.^36, 37^ After reconstituting the full 10-step pathway in *E. coli*, they showed that nhpSer plus the last four enzymes in the pathway led to FR-900098 production.^36, 37^ This implied that nhpSer was a pathway intermediate produced by enzymes FrbA, FrbB, FrbC, FrbD, possibly FrbE, plus a transaminase that is not part of the Frb biosynthetic cluster, which can be substituted for by an *E. coli* transaminase ^36–38^ (Fig. 2A).

**Figure 2.**
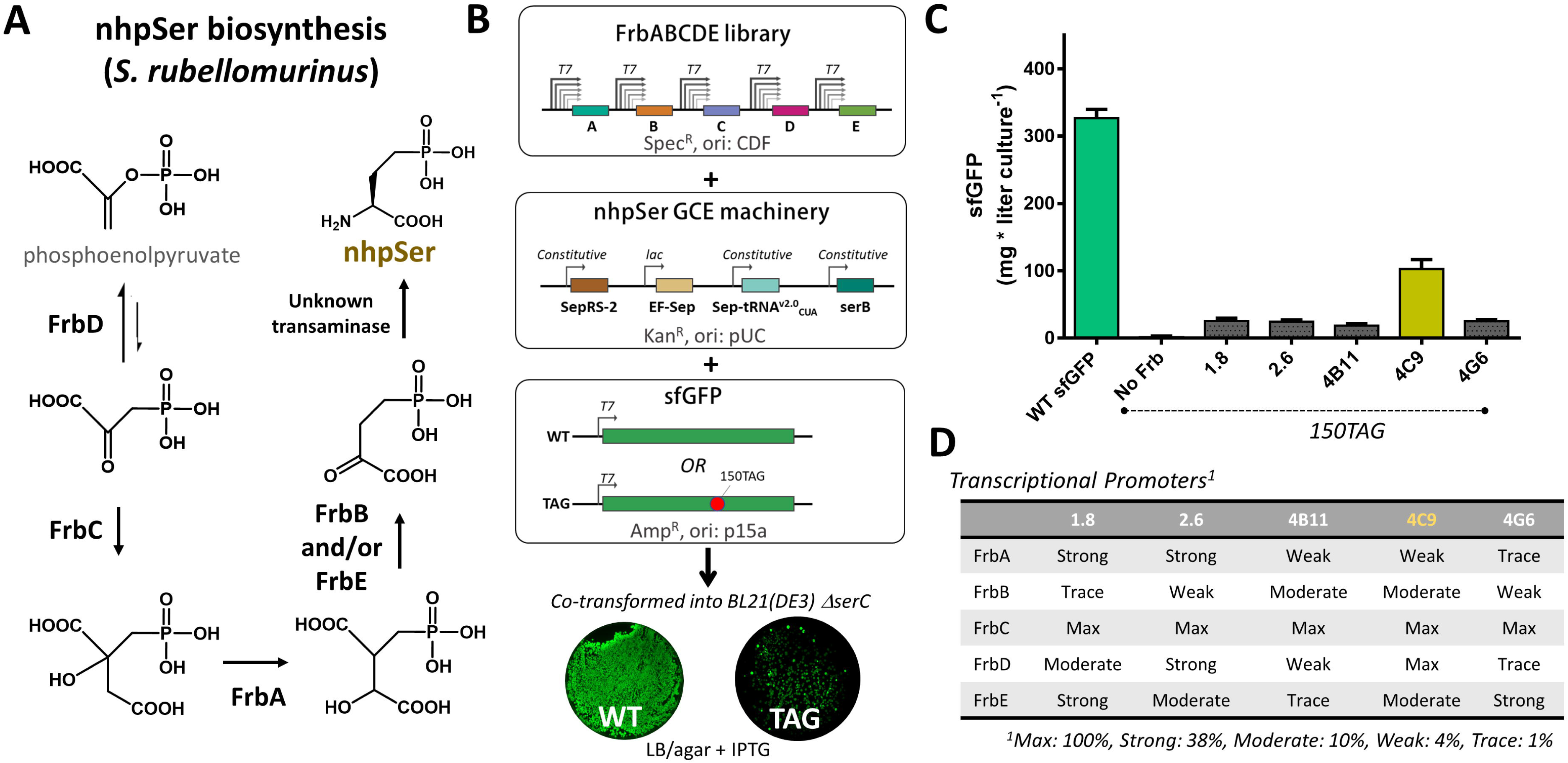
Construction of functional FrbABCDE biosynthetic pathways for translational incorporation into sfGFP in *E. coli*. (A) Biosynthetic pathway for nhpSer by *S. rubellomurinus*. The first committed step of nhpSer biosynthesis is catalyzed by a phosphoenolpyruvate mutase, FrbD, which forms the unusual C-P phosphonate bond. The energetically unfavorable equilibrium is pulled forward by the homocitrate synthase homolog FrbC via irreversible condensation of acetyl-CoA. The next two steps parallel those of the TCA cycle and are carried out by FrbA (an aconitase) and Frbs B and/or E (divergent homologs of isocitrate dehydrogenase). It is not yet known whether FrbB or FrbE catalyze this reaction, or if both are required.^36–,38^ It may be possible the FrbB and FrbE form a heterodimer, or that the two-step isocitrate dehydrogenase reaction is decoupled such that one catalyzes NADP-dependent oxidation while the other performs decarboxylation. (B) A library of FrbABCDE pathways containing all combinations of Frb enzyme transcriptional strengths was co-transformed with a plasmid housing all nhpSer GCE components and either a sfGFP-WT or sfGFP-150TAG reporter plasmid into BL21(DE3) Δ*serC* cells. (C) Comparison of nhpSer incorporation efficiency into sfGFP-150TAG for the top 5 isolated FrbABCDE pathways compared to wild-type sfGFP expression. Error bars represent standard deviations of expressions performed in triplicate. (D) Promoter identities for the Frb enzymes for each isolated pathway shown in panel C. Promoter sequences are indicated in Supplemental Fig. 1.

#### Engineering E. coli to make nhpSer for incorporation into proteins

Reconstitution of biosynthetic gene clusters in heterologous organisms benefit from tuning enzyme activities so as to maximize product while not exhausting cellular resources or creating toxic side effects.^39^ To do this for nhpSer biosynthesis, we adopted the strategy used to optimize the yields of FR-900098.^38^ First, a randomized library of T7 promoter variants was screened in the BL21(DE3) Δ*serC* cell line to identify five variants that drove sfGFP expression at discrete levels spanning two orders of magnitude (∼100, 40, 10, 4 and 1% that of wild-type) (Supplemental Fig. 2). Next, a combinatorial *FrbABCDE* library was created containing all permutations of the five Frb genes under the control of the five promoter variants (5^5^ = 3025 pathway assemblies, Fig. 2B).

To select for functional Frb pathway assemblies, we co-transformed the *FrbABCDE* library into BL21(DE3) Δ*serC* cells along with the nhpSer GCE machinery plasmid (including the constitutively over-expressed *serB* gene) and a plasmid expressing sfGFP-150TAG (Fig. 2B). Using sfGFP fluorescence as a readout of nhpSer-protein production, the top 5 performing Frb assemblies (Fig. 2C) represented 5 unique combinations of promoters (Fig. 2D). FrbC was transcribed at maximal levels in all selected clones, likely to more effectively “pull” the unfavorable equilibrium catalyzed by FrbD forward. As one of the expression vectors (4C9) strongly outperformed the others (Fig. 2C), we chose it for further characterization and named it “Frb-v1”.

Using the Frb-v1 reconstituted nhpSer pathway, single and double nhpSer incorporations into sfGFP yielded ∼120 and 30 mg/L culture, respectively (Fig.3A). These values are dramatically (∼40 fold) higher than when cells without Frb-v1 are supplemented with nhpSer at 2 mM concentration, are more than 50% of the expression levels achieved with optimized pSer GCE expression conditions, and correspond to about 40% and 13%, respectively, of wild-type sfGFP expression (Fig 3A).

**Figure 3.**
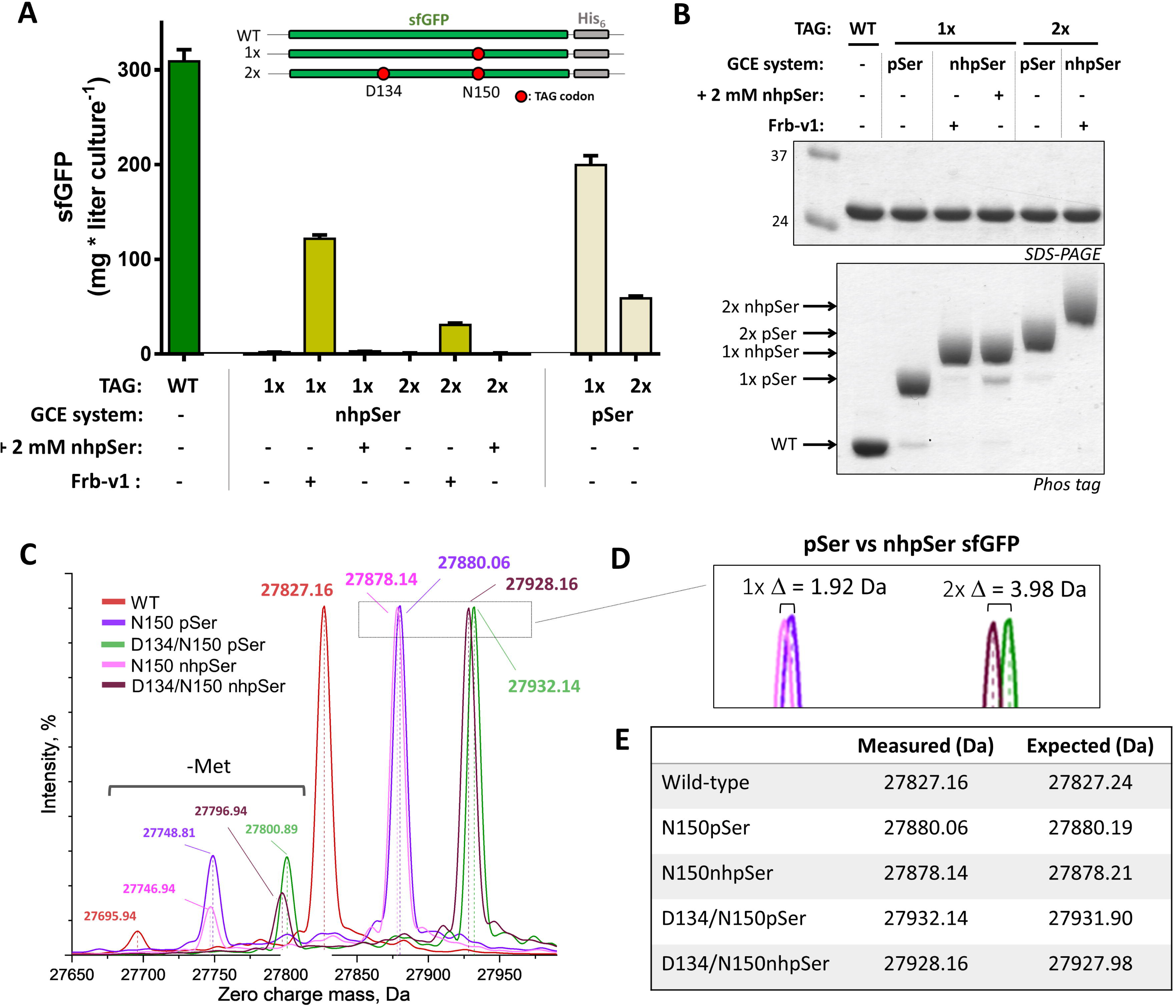
Evaluation of Frb-v1 pathway efficiency and fidelity. (A) Comparison of single (1x, 150TAG) and double site (2x, 134/150TAG) nhpSer incorporation in sfGFP when nhpSer is biosynthesized via the Frb-v1 pathway and when supplemented in the media at 2 mM concentration. Wild-type expression level and the efficiency of pSer GCE systems are shown for comparison. Error bars represent standard deviations of expressions performed in triplicate. (B) SDS-PAGE and Phos-tag gels of the purified proteins expressed in panel A. Phos-tag gels contain an acrylamide derivative that transiently interacts with phospho-groups, causing phospho-proteins to migrate slower than unmodified proteins.^35^ nhpSer-containing proteins migrate slower than their equivalent pSer containing proteins, providing a convenient method to distinguish between the two isosteres.^32, 33^ For each lane 2 μg of protein was loaded, so band intensities are not reflective of expression yield. (C) Whole-protein mass spectrometry of single and double site pSer and nhpSer incorporation into sfGFP (when nhpSer is biosynthesized by the Frb-v1 pathway). (D) Zoom in of the peaks in panel C for 1x and 2x pSer/nhpSer incorporated sfGFP confirm the expected mass differences of a bridging O to CH_2_ substitution. (E) Measured and expected whole-protein masses shown in panels C and D.

#### Fidelity assessment of nhpSer incorporation

Proteins from all the above expressions were purified by standard metal-affinity chromatography to validate nhpSer incorporation. For these purifications, C-terminal affinity tags were used to avoid co-purification of prematurely truncated peptide (Fig. 3A). All purified sfGFP forms migrated the same on SDS-PAGE gels (Fig. 3B, top), but on Phos-tag gels, the sfGFP-150nhpSer proteins produced via Frb-v1 biosynthesis and nhpSer supplementation to the media migrated identically and more slowly than sfGFP-150pSer (Fig. 3B, bottom), consistent with their reported properties^32^. Also, it appears that generating nhpSer protein using the new Frb-v1 system has lower contamination with phosphoserine sfGFP than when using 2 mM nhpSer supplementation, and also less non-phosphorylated sfGFP containing a natural amino acid at the 150-position stemming from either mis-incorporation or pSer hydrolysis. Similarly, the sfGFP-134/150nhpSer protein produced via Frb-v1 biosynthesis migrated slower than the equivalent protein doubly substituted with pSer and gave no evidence of contaminating pSer-containing protein. Electrophoretic comparison with sfGFP-134/150nhpSer made via nhpSer supplementation was not feasible due to insufficient expression levels.

We next used ultra-high resolution mass spectrometry to directly demonstrate the translational incorporation of nhpSer into sfGFP. All proteins - wild type, pSer and nhpSer sfGFP (single and double site incorporation)– had average molecular masses within 0.25 Da (<10 ppm) of their expected values (Fig. 3C-E). Evaluating the pSer-containing sfGFP side-by-side with the nhpSer-equivalent proteins confirmed that our whole-protein MS methods were sufficiently accurate to observe the small mass differences between pSer and nhpSer (expected Δ=1.98 and 3.96 Da for the O to CH_2_ substitution at the bridging γ-atom for single and double site incorporation, respectively, Fig. 3D). Further, electron transfer dissociation MS/MS fragmentation of tryptic digests unambiguously confirmed the location of nhpSer and pSer at the targeted sites D134 and N150 in the doubly modified sfGFP variants (Supplemental Figs. 3 and 4). We also did not find, within limits of detection, pSer or any natural amino acid mis-incorporated at the D134 and N150 sites of nhpSer incorporation when we expanded the search space to include all possible variations.

#### nhpSer in proteins made with Frb-v1 are resistant to hydrolysis

To confirm nhpSer-containing proteins made via the Frb-v1 pathway are indeed resistant to phosphatase activity, we incorporated pSer and nhpSer at the functionally relevant site of Ser16 of the small heat shock protein B6 (HSPB6)^40^ expressed as a peptide (residues 11-20) fusion with a highly soluble SUMO protein (Supplemental Fig. 5A). We verified nhpSer and pSer incorporation by Phos-tag gels, and upon incubation with λ-phosphatase, the pSer-containing protein was hydrolyzed while the nhpSer containing protein was not (Supplemental Fig. 5B). Collectively, these data unambiguously demonstrate that reconstitution of the FrbABCDE pathway in *E. coli* cells in combination with the nhpSer GCE machinery creates a 21-amino acid, fully autonomous organism able to efficiently synthesize protein with <u>perma</u>nent <u>phos</u>pho<u>ser</u>ine mimics site-specifically installed. We refer to this technology as “PermaPhos^Ser^”.

### Assessing nhpSer mimicry of pSer function

Development of this efficient and scalable PermaPhos^Ser^ technology, in conjunction with already established pSer GCE machinery,^7, 8^ opens the door to more easily evaluate nhpSer mimicry of pSer in a variety of contexts. To address this, we first directly measured the pKa2 values of nhpSer and pSer on a protein to assess the extent to which its charge state could influence its ability to mimic pSer at physiologic pH. Then, for each of three major regulatory outcomes of serine phosphorylation – (i) stabilization of protein-protein interactions, (ii) disruption of protein-protein interactions, and (iii) alteration of enzymatic activity – we selected one biologically relevant system for which Asp/Glu mutations appear to be poor mimetics and directly compared the impact of nhpSer *vs.* pSer *vs*. Asp/Glu.

#### Charge state of nhpSer at physiological pH

Methylene phosphonates like nhpSer have reported pKa2 values of ∼7-8, compared for ∼5-6 for phosphates,^5^ leading to concerns about the ability of nhpSer to mimic pSer. However, we found no studies that directly measured the pKa2 value of nhpSer in a protein. To directly measure their pKa2, we used the above-described SUMO-HSPB6 fusion protein with pSer and nhpSer at Ser16 (Fig. 4A) and followed the ^31^P NMR chemical shifts as a function of pH^41, 42^. We observed a well-defined ^31^P peak for nhpSer and pSer containing proteins, and consistent with previous work, increasing pH resulted in an upfield ^31^P chemical shift for nhpSer and a downfield shift for pSer^41^, with the inflection points indicating a pKa2 of 7.00 ± 0.05 for nhpSer and 5.78 ± 0.06 for pSer (Fig. 4B, Supplemental Fig. 6). This pKa2 of nhpSer is similar to the value of 7.08 seen for a phosphonate analog of phospho-lysine^41^. These data imply that at pH 7.4, nhpSer in this context exhibits an average charge of -1.7 (∼71% dianonic), compared to -2.0 for pSer, whereas the carboxylate of Asp/Glu is -1.0.

**Figure 4.**
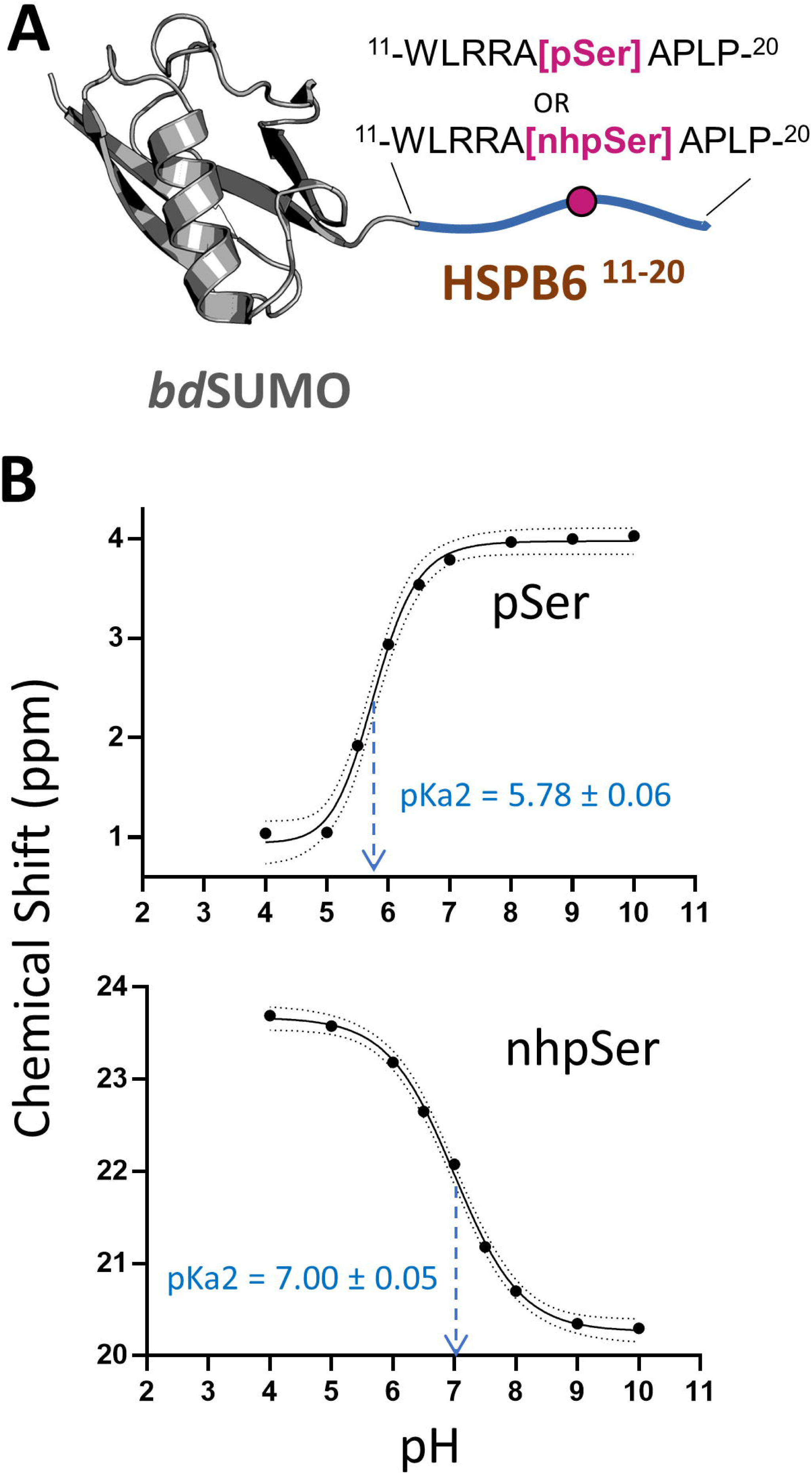
pKa2 measurement of pSer and nhpSer in context of a protein. (A) pSer and nhpSer were incorporated into site Ser16 of HSPB6 (residues 11-20) fused to a *bd*SUMO domain. This construct was chosen for NMR analysis because it was soluble at millimolar concentrations and is small, but also because the phospho-site is within an unstructured segment of the protein so that phosphorus resonance changes reflect protonation state rather than changes in tertiary or secondary structure. Phos tag gels confirming pSer and nhpSer incorporation are shown in Supplemental Fig. 5. (B) Plotting of ^31^P NMR chemical shifts revealed a sigmoidal curve from which the pKa2 of each phospho-moiety was extrapolated at the inflection point. Dotted lines represent the 95% confidence interval of the fitted curves. Raw ^31^P NMR spectra are shown in Supplemental Fig. 6.

#### Mimicking pSer-dependent stabilization of protein:protein interactions

Next, we tested the ability of nhpSer to promote pSer-dependent protein-protein interactions. For this we chose the well-studied 14-3-3 family of dimeric hub proteins that bind to and regulate hundreds of “client” pSer/pThr-containing proteins^43^ and for which it is well documented that Asp/Glu are not functional mimics.^22, 44, 45^ For a model client of 14-3-3, we chose full-length HSPB6 because it forms a structurally and thermodynamically well-characterized high-affinity complex with 14-3-3 when phosphorylated at Ser16.^40, 46^ For initial assessment of 14-3-3/client complexation, we adopted our previously reported *E. coli* expression system^8^ in which untagged 14-3-3ζ is co-expressed with His-tagged full-length HSPB6, so that 14-3-3ζ would co-purify with HSPB6 only if they formed a stable complex (Fig. 5A).

**Figure 5.**
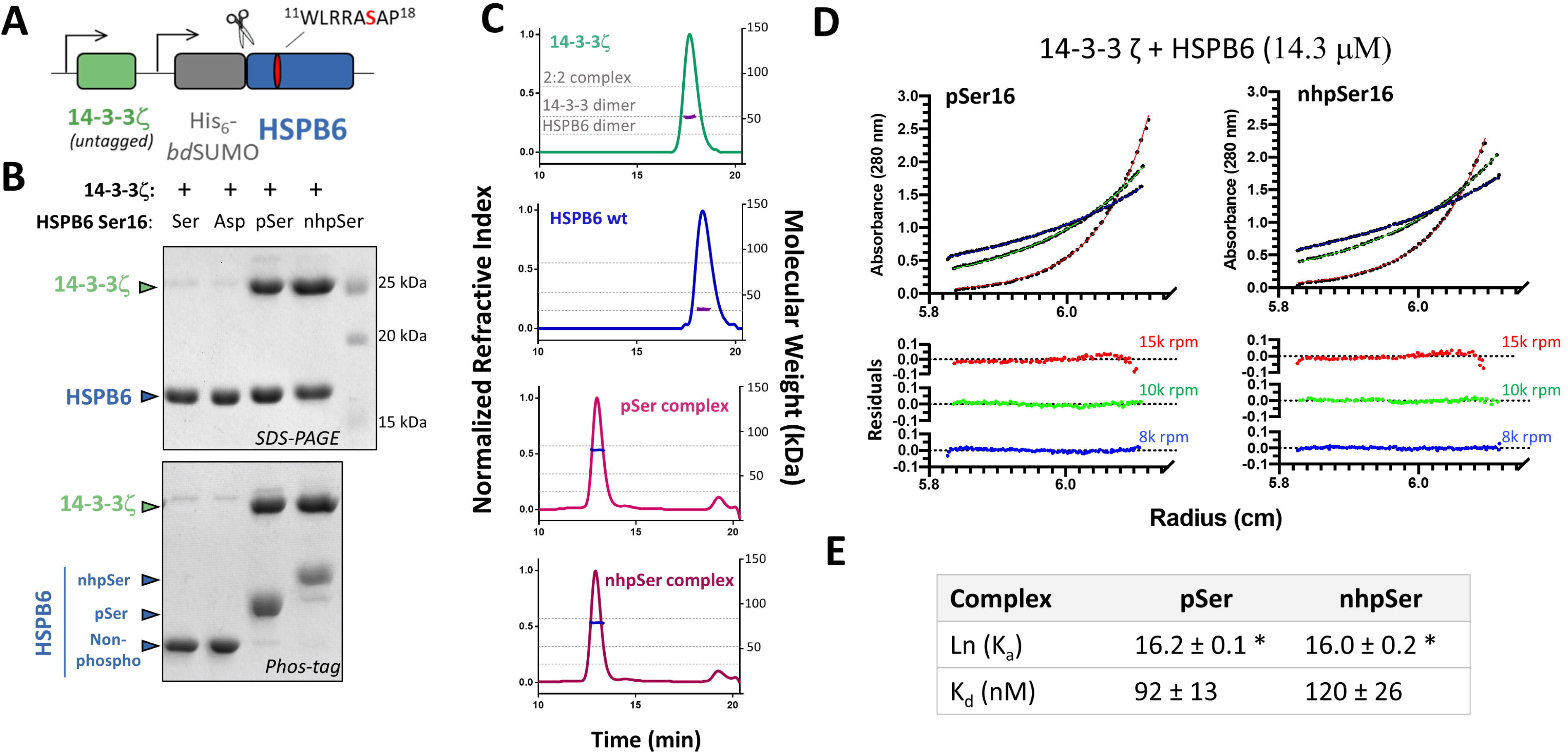
Mimicking pSer-dependent stabilization of 14-3-3/HSPB6 interactions with nhpSer. (A) Untagged 14-3-3 and His_6_-*bd*SUMO-HSPB6 proteins were co-expressed such that if they formed a tight complex, 14-3-3 co-purifies with HSPB6. At site 16 of HSPB6, which lies within a conical and well-established 14-3-3 binding site, either serine, aspartate, pSer or nhpSer were incorporated. (B) SDS-PAGE (top panel) of the purified proteins show that 14-3-3 co-purifies with HSPB6 when pSer and nhpSer, but not Ser or Asp, were incorporated at site S16. Phos-tag gel electrophoresis (bottom panel) of these proteins confirmed nhpSer incorporation into HSPB6 as indicated by it having the slowest electrophoretic mobility, with pSer being intermediate between nhpSer and wild-type or S16D HSPB6. (C) SEC-MALS of the purified protein complexes confirm 2:2 stoichiometry. Dotted gray lines denote the theoretical molecular weights of the 2:2 complex (85.4 kDa), the 14-3-3 dimer (52.4 kDa) and the HSPB6 dimer (33.0 kDa). Data statistics are shown in Supplemental Table 1. (D) Equilibrium AUC of the pSer (left) and nhpSer (right) HSPB6/14-3-3 complexes at 14.3 μM concentration at 15k (red), 10k (green) and 8k (blue) rpm. Residuals from fitting the equilibrium data to a two-state A+B ↔ AB model are shown in the bottom panels. Equilibrium data for 7.2 μM and 3.6 μM concentrations are shown in Supplemental Fig. 9. (E) Global fitting of 14-3-3/HSPB6-pSer16 and 14-3-3/HSPB6-nhpSer16 equilibria at three different protein concentrations and at three different centrifugation speeds revealed dissociation constants of 92 ± 13 and 120 ± 26 nM, respectively. Error represents 95% confidence interval range. * Denotes statistically indistinguishable calculated values by student’s unpaired t-test, *p* = 0.05; t = 0.832.

Expressing full-length HSPB6 with either Ser, pSer, nhpSer or Asp at position 16, 14-3-3ζ co-purified with HSPB6-pSer16 and with HSPB6-nhpSer16, but not HSPB6-WT or HSPB6-S16D (Fig. 5B, Supplemental Fig. 7), confirming that only pSer and nhpSer promote a stable complex between HSPB6 and 14-3-3. We further confirmed by size-exclusion chromatography coupled to multi-angle light scattering (SEC-MALS) that the complexes of 14-3-3 with HSPB6-pSer16 and with HSPB6-nhpSer16 both have 2:2 stoichiometry (Fig. 5C, Supplemental Table 1) as expected.^40^ In contrast, when either purified HSPB6-WT or the S16D variant were mixed with 14-3-3ζ, they remain un-complexed by SEC-MALS analysis (Supplemental Information Fig. 8).

We next used equilibrium analytical ultracentrifugation (AUC) to directly measure the affinities of the pSer and nhpSer promoted HSPB6/14-3-3 complexes. Global fitting of the 14-3-3/HSPB6-pSer16 and -nhpSer16 2:2 complexes fit well to a two-state model and yielded statistically indistinguishable dissociation constants of 92 ± 13 and 120 ± 26 nM, respectively (Fig. 5D and 5E, Supplemental Figs. 9 and 10). These measured K_d_ values are smaller than but consistent with the previously reported K_d_ of 560 ± 200 nM for the pSer-dependent complex extrapolated from anisotropy measurements of fluorescently labeled HSPB6.^40^ Regardless, HSPB6 with nhpSer or pSer at site S16 bind 14-3-3 with indistinguishable affinity, while the S16D variant forms no detectable complex with 14-3-3.

#### Mimicking pSer-dependent disruption of protein-protein complexes

The dimeric status of 14-3-3 isoforms ζ, ε, γ, η and β is regulated by sphingosine-dependent kinase phosphorylation of a serine – Ser58 in 14-3-3ζ – at the dimer interface (Fig. 6A)^47^. Asp/Glu phosphomimetic mutations are reported to weaken the 14-3-3 dimer, albeit poorly compared to phosphorylation^48, 49^. In the absence of methods to produce homogeneously phosphorylated 14-3-3 at this serine, characterizations of monomeric 14-3-3 have relied on additional interface mutations for full monomerization.^49^ Using pSer and PermaPhos^Ser^ GCE, we expressed and purified homogenous 14-3-3ζ with Ser, pSer, nhpSer, as well as Glu at position 58 (Fig. 6B), and evaluated their oligomeric status using SEC-MALS (Fig. 6C, Supplemental Table 2). As expected, 14-3-3ζ-WT eluted as a single dimeric peak at 50 μM initial concentration, as did the phosphomimetic S58E variant, consistent with previous work.^49^ In contrast, both 14-3-3ζ-pSer58 and 14-3-3ζ-nhpSer58 eluted exclusively as a monomer. Thus, nhpSer at position 58 recapitulates the phosphorylation-dependent monomerization of 14-3-3, while Glu at this position does not, perhaps because 14-3-3 monomerization is sustained by pSer-dependent conformational changes not imparted by a carboxylate group.

**Figure 6.**
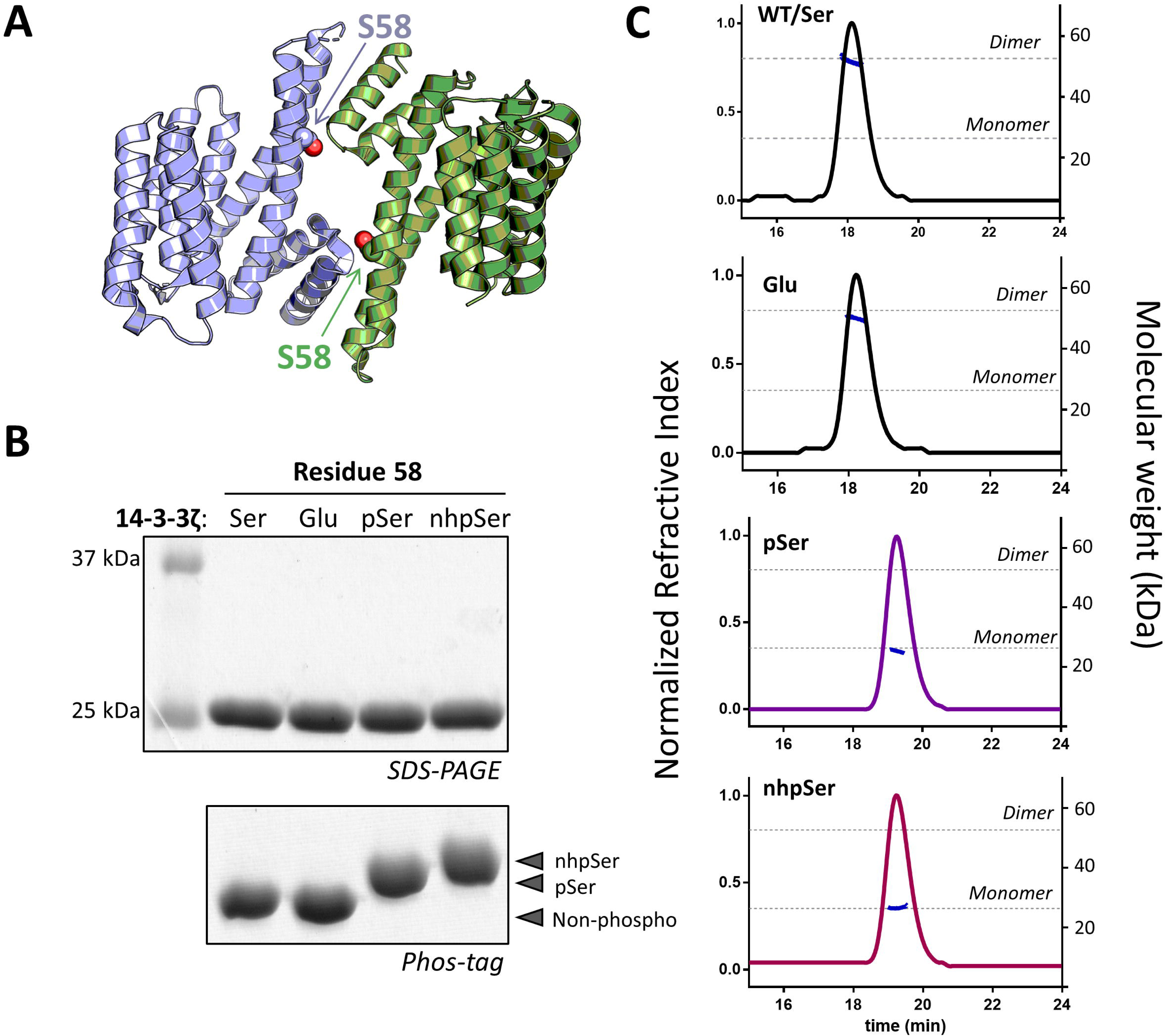
Mimicking pSer-dependent disruption of the 14-3-3 dimer. (A) Ser58 lies at the dimer interface of 14-3-3ζ (PDB 6F08). Individual protomers are colored in blue and green. (B) SDS-PAGE and Phos-tag gels confirm pSer and nhpSer incorporation into 14-3-3ζ. (C) SEC-MALS of the four 14-3-3ζ protein variants shown in panel B confirm the dimeric status of WT and S58E 14-3-3ζ, while pSer58 and nhpSer58 variants elute exclusively as a monomer. Dotted lines indicate theoretical masses of the dimeric (52.4 kDa) and monomeric 14-3-3 (26.2 kDa). All proteins were injected onto the SEC column at 50 μM initial concentration. Data statistics are shown in Supplemental Table 2.

#### Mimicking pSer-dependent activity of Glycogen Synthase Kinase-3β (GSK3β)

GSK3β is a serine/threonine kinase that regulates numerous signaling pathways. Rather than being activated by direct phosphorylation, GSK3β activity is “primed” by substrates that are Ser/Thr phosphorylated by another kinase; GSK3β then phosphorylates Ser/Thr residues four positions N-terminal to the substrate priming phospho-site (Fig. 7B).^50^ The binding of primed substrates activates GSK3β by inducing conformational changes similar to those triggered in canonical kinases by the phosphorylation of their activation loop (Supplemental Fig. 11),^51^ and simultaneously places the -4 Ser/Thr substrate residue into the activated catalytic pocket. A hallmark feature of GSK3β is the so-called “zippering” effect in which each residue phosphorylated by GSK3β can serve as the next priming site for another Ser/Thr N-terminal to it, resulting in substrate poly-phosphorylation.^52^ Asp/Glu mimetics are not considered sufficient to prime GSK3β activity.^53, 54^

**Figure 7.**
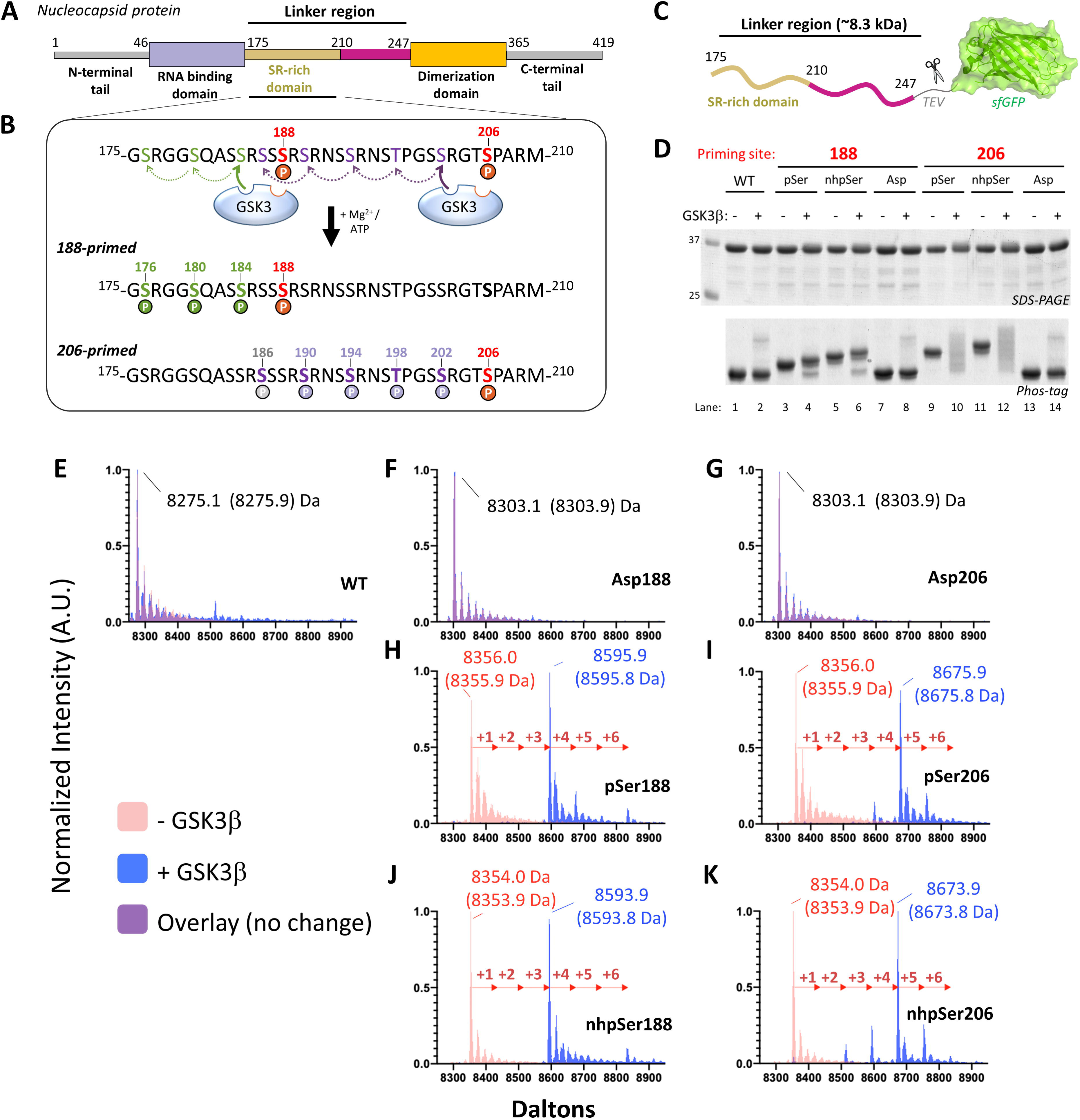
Using nhpSer as a GSK3β activator. (A) The SARS-CoV-2 Nucleocapsid Phosphoprotein (Np) contains a Serine-Arginine rich region (SR) in its linker region (Linker Np, residues 175-247) that connects the N-terminal RNA binding domain and the C-terminal dimerization domain. (B) Proposed mechanism of GSK3β poly-phosphorylation of the SR region of Np: sites S188 and S206 act as priming sites GSK3β activity when phosphorylated, resulting in two distinct tracts containing up to 3 and 5 additional sites of phosphorylation, respectively. (C) The Linker-Np was genetically fused to a TEV cleavable C-terminal sfGFP protein for enhanced solubility. (D) Ser, Asp, pSer and nhpSer were incorporated at sites S188 and S206 within the Linker-Np, purified, and mixed with ATP, Mg^2+^, and GSK3β. SDS-PAGE and Phos-tag gels of reaction products confirm phosphorylation of Linker-Np by GSK3β only in the pSer and nhpSer primed Linker-Np proteins, and not the Ser or Asp proteins, as evidenced by the gel-shifts observed in the Phos-tag gel. (E-K) Whole-protein mass Spectrometry of Linker-Np (without sfGFP) before (pink) and after (blue) GSK3β reactions. Overlapping spectra where no change occurs combine to be purple in color. (E) WT, (F) S188Asp, (G) S206Asp, (H) S188pSer, (I) 206pSer, (J) 188nhpSer, (K) 206nhpSer. Arrows indicate +80 Da increments corresponding to the addition of a phosphate group. Theoretical masses of the dominant protein species are shown in parathesis.

As a stringent test of pSer mimicry, we asked whether nhpSer could serve as a priming site to promote GSK3β activity. Due to its relevance to the current Covid-19 pandemic, we focused on the GSK3β hyperphosphorylation of the “Linker” region of the SARS-CoV-2 nucleocapsid protein (Np) (Fig. 7A). This modification is essential to the viral life cycle^55^, apparently facilitating the release of genomic RNA from Np for viral protein translation.^56–58^ Sites S188 and S206 of Np are proposed priming sites for GSK3β phosphorylation, resulting in the potential addition of 3 and 5 phosphorylation events, respectively (Fig. 7B). We expressed and purified the relevant Linker region of Np (Fig. 7C) with Ser (wild-type), pSer, nhpSer and Asp at sites 188 or 206, and incubated them with GSK3β, ATP and Mg^2+^. Phos-tag gel analysis showed that only trace amounts of wild-type, S188D and S206D variants were phosphorylated (Fig. 7D, lanes 1/2, 7/8 and 13/14), whereas Linker Np-pSer188 and -pSer206 both underwent substantial electrophoretic band shifts, indicative of phosphorylation by GSK3β (Fig. 7D, lanes 3/4 and 9/10). Analogous band shifts were observed for the Linker Np-nhpSer188 and Np-nhpSer206 variants upon incubation with GSK3β, though slightly upshifted as expected for nhpSer containing proteins (Fig. 7D, lanes 5/6 and 11/12). We also conducted the same GSK3β reactions using full-length Np and observed the same pattern of Phos-tag electrophoretic mobility shifts when Ser, Asp, pSer and nhpSer were placed at sites 188 or 206 (Supplemental Fig. 12).

We then used mass spectrometry to compare in greater detail the identity of the products resulting from pSer-primed *vs.* nhpSer-primed GSK3β phosphorylation of the Np Linker (after removing the sfGFP). This analysis showed the production of only a trace amount of phosphorylated product for wild-type, S188D and S206D (Fig. 7E-G, Supplemental Figs. 13, 14, 19), consistent with the Phos-tag gel results (Fig. 7D). However, GSK3β incubation with the pSer188 and pSer206 primed Np Linkers produced a dominant species with 3 and 4 added phospho-sites, respectively, along with other minor species (Fig. 7H and 7I). MS/MS of the dominant poly-phosphorylated species confirmed the locations of the added phosphates were exactly as expected for the GSK3β “zippering” effect (Supplemental Figs. 15, 16, 20, 21). Finally, analyses of GSK3β reacted nhpSer188 and nhpSer206 Linker-Np produced the same dominant species as the pSer primed proteins (Fig. 7J and 7K), with identical locations of the added phospho-sites (Supplemental Figs. 17, 18, 22, 23). The nhpSer206 primed Np-linker had a small population of protein with only 2 phospho-sites added instead of 4, which was not observed with pSer206 priming (compare Figs. 7I and 7K), perhaps due to slightly slower reaction kinetics. Collectively, these results show nhpSer serves as an effective functional mimic to prime GSK3-dependent phosphorylation, whereas the traditional mimetic Asp fails to do so.

## DISCUSSION

### Advantages of PermaPhos^Ser^

Though strategies have already been established to incorporate non-hydrolyzable mimics of phosphoserine into proteins, the PermaPhos^Ser^ technology we present here provides several significant advantages:

- *Translational incorporation.* First, PermaPhos^Ser^ leverages GCE technologies that enable nhpSer to be incorporated into any site of a protein irrespective of its location, including termini, structured or disordered segments, or within a protein complex. In contrast, chemical ligation techniques that are restricted in the placement of phospho-amino acid and are often time and resource intensive.
- *Increased efficiency and fidelity.* Second, PermaPhos^Ser^ employs a functional biosynthetic pathway to produce free nhpSer amino acid from the central metabolite phosphoenolpyruvate. In doing so, these cells are fully autonomous expression hosts with a 21-amino acid genetic code requiring no addition of exogenous precursor compounds or non-natural metabolites. The goal of this strategy was to overcome the reliance on media supplementation of chemically synthesized nhpSer which, as a charged molecule, does not readily cross cellular membranes restricting bioavailability and lowering overall efficiency and fidelity of protein synthesis.^59^ Indeed, only until very recently (while this manuscript was under preparation) was the first report outside the originating lab published describing successful incorporation of nhpSer via GCE to study a protein, and that success required media to be supplemented with 8 mM nhpSer to produce sufficient quantities of ubiquitin and NEDD8 proteins, and even that amount of nhpSer was still insufficient to out-compete undesired incorporation of pSer.^33^ Compared with that system, PermaPhos^Ser^ affords a ∼40-fold improvement in overall production of a test protein with increased fidelity, and enables multi-site incorporation of nhpSer. And in further tests, we demonstrate faithful and facile expression of several biologically relevant proteins and a phosphorylation-dependent protein complex in sufficient quantities for biochemical and biophysical characterization.
- *Scalability and versatility.* Third, nhpSer biosynthesis overcomes the high cost of supplementing media with chemically synthesized nhpSer, which limits the ability to produce target proteins at scale. This reliance of supplementing media with non-canonical amino acids (ncAAs) has been called the “Achilles’ heel” of GCE preventing widespread industrial adaptation of GCE technologies.^60^ Recently there have been a number of biosynthetic pathways adapted for GCE to overcome this bottleneck, though many still rely on supplementation of precursor molecules not endogenous to the expression host.^60, 61^ NhpSer biosynthesis begins from phosphoenolpyruvate, and so PermaPhos^Ser^ expression is functional in standard complex media (e.g. 2xYT and Terrific Broth). As databases of biosynthetic pathways for non-proteinogenic amino acids continue to expand,^62^ combined with easy access to synthetic genes and improved methods for heterologous pathway assemblies, PermaPhos technologies could be developed for other phospho-amino acid derivatives. While the Frb-v1 pathway has more biosynthetic steps than previous pathways assimilated for a GCE translational system, the methodology we adapted from Zhao and colleagues^38^ to optimize the Frb pathway implementation made the process relatively straightforward, and the approach may prove generally useful for incorporating other complex ncAA biosynthetic pathways.

### NhpSer as a mimic for pSer

The practical utility of PermaPhos^Ser^ technology for answering biological questions depends on how well nhpSer can mimic the functional effects of pSer. Common concerns about its mimicry are its elevated pKa2, as well as the loss the hydrogen bond-accepting oxygen. The wide range of pKa2 values reported for phosphonates prompted us to measure this directly, and given the measured pKa2 ∼ 7.0, a majority of nhpSer molecules are expected to be doubly deprotonated at physiologic pH, though this of course may vary depending on protein context. To address the second concern, we surveyed the PDB using HBPlus^63^ to assess how commonly the bridging γ-oxygen of pSer acts as a hydrogen bond acceptor in pSer dependent protein interactions. Our analysis revealed that of the 171 unique structurally-known proteins (<75% sequence identity) with pSer, the γ-oxygen of pSer was involved in a putative hydrogen bond with a protein atom in only 35 of them (∼20%). Major pSer binding protein families 14-3-3, BRCA1 C-terminal (BRCT) and WW domains do not use the γ-oxygen of pSer as a hydrogen bond acceptor to stabilize pSer-dependent interactions (Supplemental Fig. 24). Of the remaining 136 proteins, 36 contained a pSer moiety for which water was modeled in a position to act as potential hydrogen bond donor to the γ-oxygen of pSer. These observations show that the lack of hydrogen bonding ability of the bridging CH_2_ does not inherently disqualify the potential of nhpSer to recapitulate pSer-dependent interactions.

Consequently, we carried out an extensive side-by-side characterization of nhpSer mimicry for pSer with three stringent test cases: 14-3-3/client phosphoserine-dependent complexation, pSer-dependent monomerization of 14-3-3, and the priming of GSK3β kinase activity. Hydrogen bonding analyses of 14-3-3 complexes (Supplemental Fig. 24A) and GSK3 with substrate peptide (Supplemental Fig. 11B) indicated that in these cases the γ-oxygens of pSer were not involved in complex stabilization. No structure of phosphorylated monomeric 14-3-3 has been solved and so whether the pSer γ-oxygen is involved in interactions is unknown. Nevertheless, for all three cases, our results confirm that nhpSer was able to mimic the effects of serine phosphorylation under the conditions tested, while Asp/Glu failed in all cases to mimic these changes. While noteworthy differences were not detected in these diverse characterizations, we have no doubt that nhpSer will not be a perfect mimic of pSer in all contexts. Indeed, in the recent elegant study by Stuber *et al*.,^33^ while a nhpSer65-variant of ubiquitin did adopt the functionally important pSer- dependent conformational state that was not adopted by the Asp/Glu variants, it did so with lower occupancy than authentically phosphorylated ubiquitin. In this case, the difference in nhpSer and pSer charge states was credited for the disparity, however our pKa2 measurement of nhpSer would suggest the large majority of nhpSer should still be dianionic like pSer, and so additional mechanisms may be at play to explain the discrepancy. Nevertheless, it is very hard to imagine any context in which Asp/Glu would afford a more effective mimic than nhpSer.

### Outlook

Collectively, the PermaPhos^Ser^ system with the Frb-v1 pathway provides an efficient method for producing permanently phosphorylated proteins in *E. coli*. Its advent begins a new era in which the use of Asp/Glu as stable, phosphatase resistant phosphomimetics can be phased out and replaced with autonomous incorporation of nhpSer. The combination of PermaPhos^Ser^ and previously developed, efficient pSer GCE systems together open the door to much more informative biochemical and biophysical studies of wide range of phosphoproteins, as well as the development of tools and therapeutics that target kinases, phosphatases, and other phospho-dependent protein systems. Moving forward, while the metabolic levels of nhpSer produced by Frb-v1 are sufficient for use, improvements could be possible. For instance, the Frb- v1 vector may not represent the best possible implementation of the pathway, because we only varied the transcriptional levels of the five enzymes using inducible promoters. Identifying the unknown transaminase, and regulating its expression levels could prove advantageous, as would testing the activity of other FrbABCDE-like orthologs or creating fusions that anchor Frb enzymes near each other. As another potential development, it should be possible to create mammalian cells with the ability to autonomously synthesize specific, permanently phosphorylated proteins, because all components of the nhpSer GCE machinery are compatible with mammalian expression hosts. It is reasonable to speculate that such a system would provide the practical advantages for nhpSer protein production as shown here for *E. coli*, especially when considering the existing mammalian nhpSer GCE system requires 25 mM nhpSer added to the media for function, and still the majority of target protein has natural amino acid at the intended site of incorporation^32^.

## METHODS

### Strains

BL21(DE3) Δ*serC* and BL21(DE3) Δ*serB* were gifts from Jason Chin and Jesse Rinehart, respectively. B95(DE3) Δ*A* Δ*fabR* Δ*serB* was generated at the OSU Unnatural Protein Facility as previously described.^8^ DH10b was purchased from Thermo Fisher Scientific. The PPY strain used to generate SLiCE^64^ cloning extract was a gift from Yongwei Zhang (Albert Einstein College of Medicine).

### Molecular biology reagents

Oligonucleotide primers and double stranded DNA fragments were synthesized by Integrated DNA Technologies (Coralville, IA). Molecular biology reagents including restriction enzymes, T4 ligase and polymerases were purchased either from Thermo Fisher Scientific or New England Biolabs. DNA Miniprep, Midiprep, PCR cleanup and gel extraction kits were purchased from Machery Nagel.

### Generating the Frb-v1 nhpSer biosynthetic pathway

#### Selection of attenuated T7 promoters

A library of mutant T7 promoters with altered transcriptional activity was generated and placed in front of a fluorescent reporter sfGFP gene. Mutant promoters with targeted transcriptional efficiency were selected in a similar manner as previously described, except that the expression strain was BL21(DE3) Δ*serC*.^38^

#### FrbABCDE library assembly

The FrbABCDE library assembly strategy was adapted Zhao and colleagues^38^ such that all permutations of attenutated transcriptional promoters in front of each of the five Frb genes were made via Golden Gate assembly (5^5^ = 3125 combinations). Each gene was flanked by its own promoter and terminator so that each is transcribed independently. The plasmid backbone contained a CDF origin of replication and spectinomycin resistance to ensure compatibility and stable propagation with the nhpSer GCE machinery (pUC origin/kanamycin resistance) and target protein expression plasmids (p15a origin/ampicillin resistance).

#### Screening for functional FrbABCDE assemblies

The pCDF-FrbABCDE_lib was screened for assemblies able to synthesize nhpSer for translational incorporation into sfGFP-150TAG. To do this, the pCDF-FrbABCDE_lib was electroporated with the nhpSer GCE machinery plasmid, pSF-nhpSer and the reporter expression plasmid, pRBC-sfGFP-150TAG, into BL21(DE3) Δ*serC* cells. Cells were plated on inducing LB/agar plates, and 92 top fluorescent clones were selected. Plasmid DNA the top 6 clones were isolated and pathway functionality was re-tested in subsequent expressions at 50 mL scale, in triplicate. Promoter strengths were identified by sequencing.

### Quantification of sfGFP expression

Yield of sfGFP expressed per liter culture was calculated by measuring in-cell fluorescence of sfGFP and subtracting the contribution of cell auto-fluorescence (measured from the same density of cells not expressing any sfGFP construct). Fluorescence values were converted to mass of sfGFP per liter culture based on a standard curve of purified sfGFP. All values reported are the average of three independent replicate cultures, and error bars represent the standard deviation.

### nhpSer-protein expression

The pSF-nhpSer machinery plasmid (a kind gift from Jason Chin) was used for selective nhpSer incorporation.^32^ This plasmid expresses the SepRS-2^7^ under a constitutive GlnRS promoter, the Sep-tRNA^v2.0^_CUA_ under a constitutive lpp promoter,^34^ the EF-Sep under the control of a lac promoter,^6^ and *E. coli serB* under a strong constitutive recA promoter.

#### Expression using nhpSer media supplementation

Fresh transformations were performed for all expressions in this study. Expressions with nhpSer media supplementation were performed essentially as previously described.^7^ Briefly, BL21(DE3) Δ*serC* cells were co-transformed with pRBC-sfGFP-150TAG or pRBC-sfGFP-134/150TAG and pSF-nhpSer. Cells were grown and protein expressed in 2xYT media induced with the addition of 1 mM IPTG and 2 mM nhpSer (DL-AP4, Abcam) at 37 °C for 18 h.

#### Expression using the Frb-v1 pathway

The pCDF-Frb-v1.0, pSF-nhpSer and the appropriate pRBC plasmid were co-transformed simultaneously by electroporation into BL21(DE3) Δ*serC* cells. Expressions were performed in Terrific Broth (TB: 12 g/L tryptone, 24 g/L yeast extract, 0.5% glycerol, 72 mM K_2_HPO_4_, 17 mM KH_2_PO_4_) and initiated by the addition of 0.5 mM IPTG once cells reached an OD_600_ ∼1.0. Cultures were harvested 24 hrs after the addition of IPTG.

### pSer-protein expression

The pKW2-EFSep machinery plasmid (a kind gift from Jason Chin) was used to enable pSer incorporation, which expresses SepRS-2 under a constitutive GlnRS promoter, the Sep-tRNA_CUA_ G4 under a constitutive lpp promoter, and the EF-Sep under the control of a lac promoter.^7^ Proteins with pSer were expressed from the pRBC plasmid exactly as previously described^8^ in either BL21(DE3) Δ*serB* or B95(DE3) Δ*A* Δ*fabR* Δ*serB*. The latter, which is a partially recoded derivative of BL21(DE3),^65^ lacks Release Factor-1 and therefore was used to avoid truncated protein.^8^

### Protein purification

All proteins were purified by standard metal affinity chromatography using TALON or Ni-NTA resin. sfGFP and sfGFP fusion proteins were eluted with using 300 mM imidazole, while 14-3-3ζ, HSPB6 (and their complexes with 14-3-3ζ), and full-length SARS-CoV-2 nucleocapsid protein were eluted using 30 nM untagged *bd*SENP1 on-column proteolytic cleavage of their N-terminal His_6_-*bd*SUMO tags, as previously described.^66, 67^ After elution, if needed, proteins were further purified by gel-filtration on a 10/300 Superdex S75 column (GE Healthcare).

#### λ-Phosphatase Hydrolysis

8.3 μM *bd*SUMO-HSPB6^11–20^ fusion protein containing pSer or nhpSer at site S16 of HSPB6 were reacted with 3 units of λ phosphatase (NEB) according to manufacturer’s guidelines. Reactions were quenched at the indicated time points by acetone precipitation. Precipitated proteins were pelleted at 20k rcf at 4 °C, dried and resuspended in 1x SDS sample buffer for SDS and Phos-tag gel analysis.

#### ^31^P Nuclear Magnetic Resonance (NMR)

*bd*SUMO-HSPB6^11–20^ proteins in 25 mM Tris pH 7.4, 25 mM NaCl at 5 mM concentration were diluted 10-fold in a buffer with pH ranging from 4 to 10. This dilution buffer contained 100 mM sodium acetate (pKa 4.8), 100 mM Bis-Tris (pKa 6.5), 100 mM Tris (pKa 8.1), 100 mM CHES (pKa 9.3), 25 mM NaCl, and 10% D_2_O, and was pH’d between 4 and 10. By including four reagents with pKa’s ranging from 4.8 to 9.3, buffering was effective from pH 4 to 10. Thus, all proteins for NMR analysis, regardless of pH, were in the identical buffering matrix. After dilution, any insoluble protein was precipitated by centrifugation. Samples were then transferred to 5 mm NMR tubes (Norell) and experiments were conducted on a Bruker Advance III 500 MHz NMR Spectrometer outfitted with a 5 mm BBOF probe. Data were collected with a spectral window of 99.5774 ppm, 65,536 real plus imaginary points, a D1 of 2.0 sec, and between 1,024 and 13,000 scans. All NMR experiments were collected at 25 °C. NMR data were processed (apodized, zero filled, Fourier Transformed, and phased) and analyzed in Bruker Topspin. Labeled peaks were exported to Graphpad Prism 8 and plotted against pH of the buffer; the plotted points were interpolated to a Sigmoidal, 4 PL function and the pKa2 was determined at the calculated inflection point +/- 95% Confidence Interval.

#### Sedimentation Equilibrium Analytical Ultra Centrifugation (AUC)

Protein samples for AUC were serially diluted to an Abs at 280nm of 1.0 (14.3 μM), 0.5 (7.2 μM), and 0.25 (3.6 μM) into 50 mM Tris pH 7.4, 150 mM NaCl, 1 mM TCEP. AUC experiments were performed using a Beckman Coulter Optima XL-A ultracentrifuge equipped with absorbance optics (Brea, CA). Protein-partial-specific volumes as well as buffer densities and viscosities were estimated using the software sednterp^68^. Experiments were performed using an An-60-ti rotor at three speeds: 8,000, 10,000, and 15,000 rpm all at 5 °C. Scans at a wavelength of 280 nm were taken every 3 hours to check that equilibrium had been achieved, with finals scans taken a minimum of 30 hours after reaching speed. Data were fit to a simple heterodimerization model (A+B ⇓→ AB) using fixed masses of 33031 Da for a HSPB6 dimer, and 52395 Da for a 14-3-3 ζ dimer with the Heteroanalysis software (version 1.1.60, University of Connecticut). Proteins were loaded on SDS-PAGE and Phos-tag gels before and after AUC experiments to confirm sample integrity was maintained during data acquisition. Calculated Ln Ka values for pSer and nhpSer containing complexes were compared by student’s unpaired two-tailed t-test (*p* = 0.05) to determine statistical significance.

#### Size Exclusion Chromatography coupled to Multiangle Light Scattering (SEC-MALS)

Experimental molecular weights were obtained by size exclusion chromatography (SEC) using an AKTA FPLC (GE Healthcare) coupled to a DAWN multi-angle light scattering (MALS) and Optilab refractive index system (Wyatt Technology). Size exclusions were conducted on a Superdex200 10/300 GL column (Cytiva Life Sciences) pre-equilibrated in 50 mM Tris pH 7.4, 150 mM NaCl, 0.5 mM TCEP at room temperature. Protein samples were prepared at 50 µM and injected at a flow rate of 0.8 mL/min. Duplicate datasets were analyzed using ASTRA software package, version 8 (Wyatt Technology).

#### GSK3β Kinase reactions

80 μM of target protein (Linker Np-sfGFP or full-length Np) was incubated with 80 nM of GSK3β in a buffer containing 50 mM Tris pH 7.4, 150 mM NaCl (350 mM NaCl for Linker Np-sfGFP), 10 mM MgCl_2_, and 1 mM ATP. Reactions were incubated at 37 °C for 20 hours. Reactions were then quenched for analysis by boiling in 1x SDS sample buffer and loaded on SDS-PAGE and Phos-tag gel, or frozen at -20 °C in preparation for mass spectrometry.

### Mass Spectrometry of phosphorylated sfGFP

#### Intact sfGFP

A Waters nanoAcquity UPLC system (Waters, Milford, MA) was coupled online to an Orbitrap Fusion Lumos Tribrid ETD mass spectrometer (Thermo Fisher Scientific). Protein samples were diluted 25 times in 0.1% formic acid. Proteins were loaded onto a trap 2G nanoAcquity UPLC Trap C4 Column (100mm, 50mm, 5um) at a flow rate of 5 μl/min for 5 minutes. The separation was performed on an Acquity UPLC BEH C4 column (100um, 100mm, 1.7um, 300Å). Column temperature was maintained at 37°C using the integrated column heater. Solvent A was 0.1% formic acid in LC-MS grade water and solvent B was 0.1% formic acid in LC-MS grade acetonitrile. The separation was performed at a flow rate of 0.5 μl/min, and using linear gradients of 3–10% B for 3 minutes, 10–30% B for 17 minutes, 30–90% B for 3 minutes, 90% B for 4 minutes, 95–3% B for 1 minutes, 3% B for 7 minutes. Total method length was 35 minutes. The outlet of the column was connected to the Thermo Nanospray Flex ion source and +2300V were applied to the needle.

MS acquisition method was optimized for the target sfGFPs. Optimum setting for in-source dissociation was found to be 15%. Full MS spectra were acquired over m/z 400-2000 because this mass range contained all charge states observed for the target proteins. Scans with resolution 240000 at m/z 200 with accumulation of 10 microscans were sufficient to resolve the isotopic distribution.

Raw data were deconvoluted using UniDec software^69^. Charge range and mass range was set 5 to 50 and 5kDa to 50kDa respectively. Automatic m/z peak width was determined to be ∼0.07 Th. Split Gaussian/Lorentzian was selected as peak shape function. Deisotoping was switched off. Exported deconvoluted spectra for different files were juxtaposed in OriginPro 2021 (v9.8).

#### Trypsin digestion

Protein samples were diluted 25-fold in 50 mM ammonium bicarbonate. To prevent disulfide bonds of the proteins, the samples were incubated at 56 °C for 1 hour with 5 mM dithiothreitol (ThermoFisher). Then, the samples were incubated with 10 mM iodoacetamide (MilliPore Sigma) for 1 hour at room temperature in the dark in order to carbamidomethylate cysteine residues. Samples were digested overnight at 37 °C using Trypsin Gold (Mass Spectrometry Grade, Promega). After digestion, samples were spun down at 12000 rcf for 30 s to collect condensate, and the digestion was stopped by addition of 0.5% (v/v) trifluoroacetic acid. Samples were centrifuged at 12000 rcf for 10 minutes and then transferred to LC vials.

#### Bottom-up mass spectrometry

Peptides were loaded onto a trap 2G nanoAcquity UPLC Trap C18 Column (100mm, 50mm, 5um) at a flow rate of 5 μl/min for 5 minutes. The separation was performed on a commercially available Acquity UPLC Peptide BEH C18 column (100um, 100mm, 1.7um). LC gradient and other parameters were identical to the method described above for the top-down analysis.

MS1 spectra were acquired at a resolution of 120,000 (at m/z 200) in the Orbitrap using a maximum IT of 50 ms and an automatic gain control (AGC) target value of 2E5. For MS2 spectra, up to 10 peptide precursors were isolated for fragmentation (isolation width of 1.6 Th, maximum IT of 10 ms, AGC value of 2e4). Precursors were fragmented by ETD and analyzed in the Orbitrap at a resolution of 30,000 at m/z 200. Reagent injection time and other ETD parameters were set to be selected automatically based on the calibration parameters. The dynamic exclusion duration of fragmented precursor ions was set to 30 s.

Raw files were processed in Thermo Proteome Discoverer 2.3. Precursor ion mass tolerance was set to 5 ppm, while fragment ion mass tolerance was 0.02 Da. The SequestHT search engine was used to search against the database containing target sfGFP sequence and variation databases. Variation databases were created for the incorporation sites to test for non-cognate incorporation. Additional dynamic modifications were created for aspartate and asparagine conversion into phosphoserine (+51.9714, +52.9554) and non-hydrolysable phosphoserine (+49.9922, + 50.9672). c and z ions only were considered for peptide spectrum matching. MS1 precursor quantification was used for label-free quantitation of the peptides.

### Mass Spectrometry of Linker-Np

Amicon Ultra 3,000 Da cut-off centrifugal filter units were used to buffer exchange Linker-Np samples into 200mM ammonium acetate, after which the proteins were diluted to 10µM in 15% acetonitrile and 0.1% formic acid. All samples were analyzed on an Agilent 6545XT equipped with an e-MSion ExD cell (model AQ-250) and ionized through direct infusion into a dual AJS electrospray source or sprayed with a custom static nano electrospray source. The instrument was operating in dynamic range mode (2 GHz) and spectra were collected at a rate of 1 spectrum per second. A scan range of 400-2,400 m/z was used to detect intact masses in MS1 mode and 120-2,400 m/z in MS2. The Linker-Np precursors were isolated and fragmented through a targeted acquisition run. Electron Capture Dissociation (ECD) was used to fragment the gas phase protein ions. An additional 10V of collision energy was required to fragment the multiply phosphorylated Linker-Np proteins. MS2 fragmentation spectra were acquired and averaged for approximately 6 minutes. The Agilent Mass Hunter BioConfirm B.09 software was used to deconvolute the m/z spectra and determine the intact protein masses. Fragmentation spectra were analyzed with the ExD Viewer v4.1.14. The fragment ions were compared to theoretical fragmentation data based on the known sequence of Linker-Np. Fragment ions were matched by considering the experimental mass to charge values and isotopic distribution. The maximum error allowed for mass to charge matching was 20 ppm. The experimental spectra were searched for b, y, c, and z ions, corresponding to ECD and CID-type product ions. The presence of a phosphorylation was determined by a mass shift of +79.9 relative to the theoretical mass. Non-hydrolysable phosphoserine moieties were identified by matching the theoretical methyl phosphonate fragment in position S188 or S206.

### HBplus analysis of pSer containing structures

Structures determined at < 2.5 Å resolution containing phosphoserine (residue ID: SEP) within a peptide chain were pulled from the Protein Data Bank (971 entries). The PISCES server^70^ was used to filter PDBs with <75% sequence identity, producing a list with 171 unique structures containing pSer. PDB files for these structures were downloaded and HBplus^63^ was used to assess hydrogen bonds for pSer residues in each structure using default parameters, except that the maximum donor to acceptor (D-A) distance was set to 3.5 Å.

## ACKNOWLEDGEMENTS

We thank Jason Chin for the pSF-nhpSer and pKW2-EFSep machinery plasmids. We are grateful for the mass spectrometry assistance provided by e-MSion, Inc (Corvallis, OR). We also thank Dr. Afua Nyarko for SEC-MALS facility support.

## DECLARATIONS

The authors declare that they have no conflict of interest.

## AUTHOR CONTRIBUTIONS

Project conceptualization came from P.Z., R.A.M., and R.B.C. Data acquisition and interpretation was carried out by P.Z., R.F., A.V., S.S., P.R., N.N.S. and R.B.C. The original draft was written by P.Z. and R.B.C. Review and editing of the paper was carried out by P.Z., R.B.C., P.A.K., and R.A.M. Funding was acquired by R.B.C., R.A.M. and J.S.B. Supervision of the work was responsibility of R.B.C. and R.A.M. All authors read and approved the submitted version.

