## Supplemental Information for "PermaPhos^Ser^: autonomous synthesis of functional, permanently phosphorylated proteins"

### SUPPLEMENTAL TABLES

**Supplemental Table 1.** SEC-MALS statistics for 14-3-3ζ/HSPB6 complexes shown in Main Text Figure 5C.

| | Mw (kDa) | Uncertainty | Calculated mass (µg) | Mass fraction (%) |
|---|---|---|---|---|
| WT_1433_run1 | 51.7 | 0.20% | 106.76 | 100 |
| WT_1433_run2 | 51.5 | 0.30% | 96.33 | 100 |
| Average | 51.6 | | 101.55 | 100 |
| Standard deviation | 0.1 | | 7.38 | 0 |
| % Standard deviation | 0.2 | | 7.26 | 0 |
| Minimum | 51.5 | | 96.33 | 100 |
| Maximum | 51.7 | | 106.76 | 100 |

| | Mw (kDa) | Uncertainty | Calculated mass (µg) | Mass fraction (%) |
|---|---|---|---|---|
| HSPB6 WT run 1 | 34.4 | 0.30% | 108.17 | 100 |
| HSPB6 WT run 2 | 34.5 | 0.30% | 98.02 | 100 |
| Average | 34.4 | | 103.1 | 100 |
| Standard deviation | 0.1 | | 7.18 | 0 |
| % Standard deviation | 0.3 | | 6.96 | 0 |
| Minimum | 34.4 | | 98.02 | 100 |
| Maximum | 34.5 | | 108.17 | 100 |

| | Mw (kDa) | Uncertainty | Calculated mass (µg) | Mass fraction (%) |
|---|---|---|---|---|
| pSer Complex run 1 | 79.1 | 0.90% | 393.4 | 100 |
| pSer Complex run 2 | 78.9 | 0.90% | 358.77 | 100 |
| Average | 79 | | 376.09 | 100 |
| Standard deviation | 0.1 | | 24.49 | 0 |
| % Standard deviation | 0.2 | | 6.51 | 0 |
| Minimum | 78.9 | | 358.77 | 100 |
| Maximum | 79.1 | | 393.4 | 100 |

| | Mw (kDa) | Uncertainty | Calculated mass (µg) | Mass fraction (%) |
|---|---|---|---|---|
| nhpSer Complex run 1 | 80.1 | 1.00% | 389.63 | 100 |
| nhpSer Complex run 2 | 80.2 | 1.00% | 384.83 | 100 |
| Average | 80.1 | | 387.23 | 100 |
| Standard deviation | 0 | | 3.4 | 0 |
| % Standard deviation | 0.1 | | 0.88 | 0 |
| Minimum | 80.1 | | 384.83 | 100 |
| Maximum | 80.2 | | 389.63 | 100 |

**Supplemental Table 2.** SEC-MALS statistics for 14-3-3ζ S58 variants showin in Main Text Figure 6C.

| | Mw (kDa) | Uncertainty | Calculated mass (µg) | Mass fraction (%) |
|---|---|---|---|---|
| WT_run1 | 51.4 | 1.30% | 88.82 | 96.2 |
| WT_run2 | 51.1 | 1.40% | 83.89 | 95.9 |
| Average | 51.2 | | 86.36 | 96 |
| Standard deviation | 0.3 | | 3.49 | 0.2 |
| % Standard deviation | 0.5 | | 4.04 | 0.2 |
| Minimum | 51.1 | | 83.89 | 95.9 |
| Maximum | 51.4 | | 88.82 | 96.2 |

| | Mw (kDa) | Uncertainty | Calculated mass (µg) | Mass fraction (%) |
|---|---|---|---|---|
| S58E_run1 | 50.4 | 0.20% | 95.61 | 100 |
| S58E_run2 | 50 | 0.30% | 94.32 | 100 |
| Average | 50.2 | | 94.96 | 100 |
| Standard deviation | 0.3 | | 0.92 | 0 |
| % Standard deviation | 0.5 | | 0.97 | 0 |
| Minimum | 50 | | 94.32 | 100 |
| Maximum | 50.4 | | 95.61 | 100 |

| | Mw (kDa) | Uncertainty | Calculated mass (µg) | Mass fraction (%) |
|---|---|---|---|---|
| pSer58_run1 | 25.3 | 2.00% | 85.76 | 100 |
| pSer58_run2 | 25.9 | 2.30% | 106.09 | 100 |
| Average | 25.6 | | 95.92 | 100 |
| Standard deviation | 0.5 | | 14.38 | 0 |
| % Standard deviation | 1.8 | | 14.99 | 0 |
| Minimum | 25.3 | | 85.76 | 100 |
| Maximum | 25.9 | | 106.09 | 100 |

| | Mw (kDa) | Uncertainty | Calculated mass (µg) | Mass fraction (%) |
|---|---|---|---|---|
| nhpSer58_run1 | 26.5 | 1.40% | 107.23 | 100 |
| nhpSer58_run2 | 26.6 | 1.00% | 101.02 | 100 |
| Average | 26.5 | | 104.13 | 100 |
| Standard deviation | 0 | | 4.39 | 0 |
| % Standard deviation | 0.1 | | 4.22 | 0 |
| Minimum | 26.5 | | 101.02 | 100 |
| Maximum | 26.6 | | 107.23 | 100 |

### SUPPLEMENTAL FIGURES

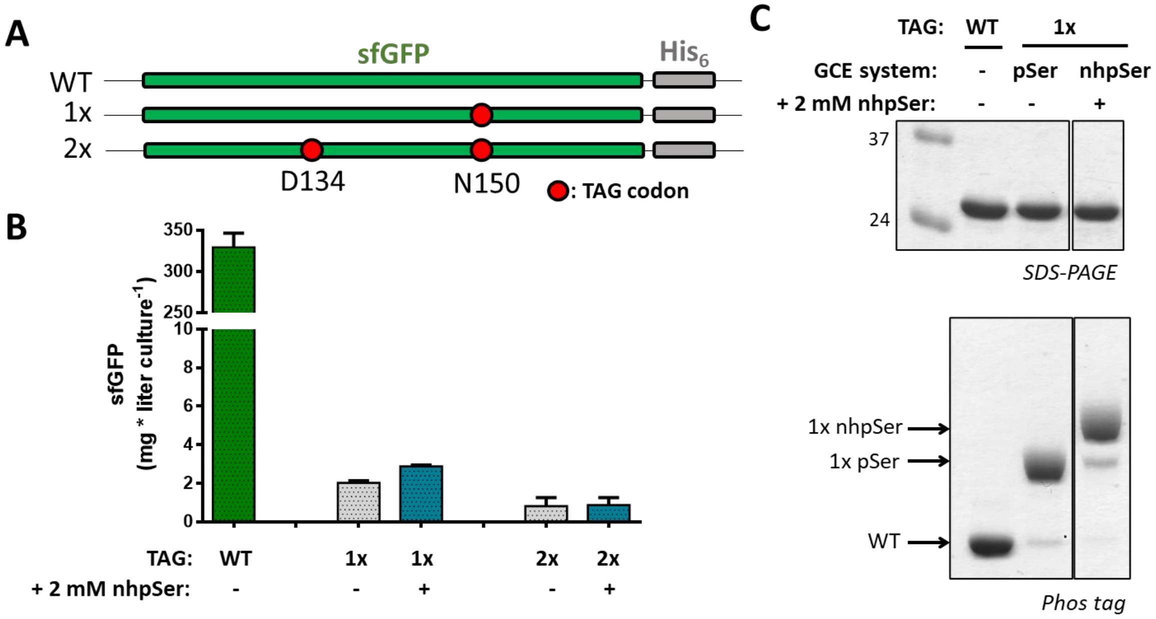


**Supplemental Figure 1**. Efficiency of nhpSer incorporation into sfGFP using the current GCE system. (A) sfGFP fluorescent reporter constructs containing one (1x, 150TAG) and two (2x, 134/150TAG) amber stop codons. (B) sfGFP expression yields based on in-cell fluorescence measurements using the current nhpSer GCE in which nhpSer is supplemented to the media at 2 mM final concentration. Expressions were performed in BL21(DE3) Δ*serC* cells using 2xYT media and 1 mM IPTG to induce expression. Cells were cultured for 18 hrs at 37 °C prior to fluorescence measurements. Yield of sfGFP per liter culture was calculated by subtracting the contribution of auto-fluorescence from cells not expressing any sfGFP construct from the fluorescence of the indicated cultures, and then fluorescence values were converted to sfGFP concentration based on a standard curve of purified sfGFP. Error bars represent standard deviations of expressions performed in triplicate. (C) SDS-PAGE and Phos tag gels of purified sfGFP wild-type, pSer150 and nhpSer150 (produced via media supplementation). The slower electrophoretic mobility of sfGFP-nhpSer150 compared to sfGFP-pSer150 is consistent with previous reports.[1] The full (uncropped) gels are shown in the main text Fig. 3B.

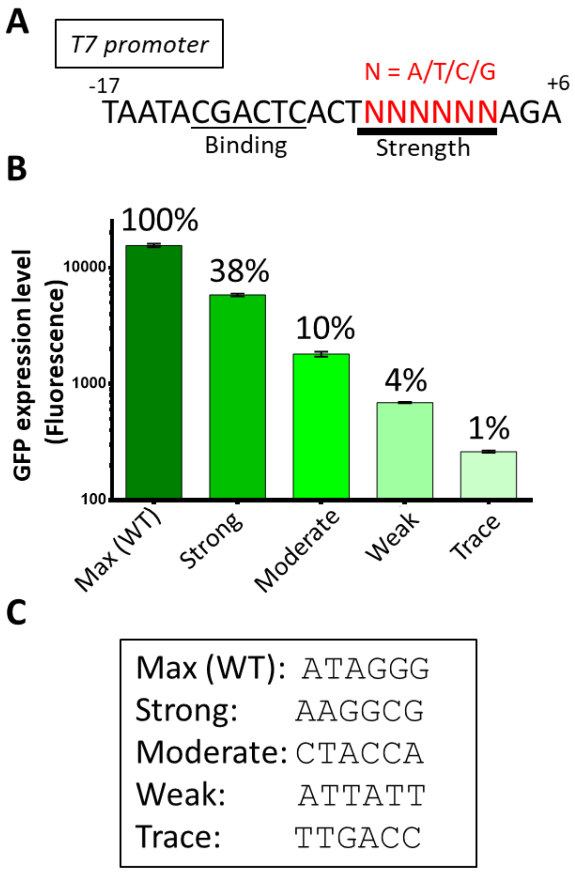


**Supplemental Figure 2**. Generating T7 promoter variants with attenuated transcriptional strengths. (A) Sequence of the T7 transcriptional promoter highlighting the region regulating transcription initiation strength.[2] Nucleotides highlighted in red were randomized to all four nucleotides and placed in front of an sfGFP reporter gene. (B) After screening the library for promoter variants conferring attenuated sfGFP expression, four were identified displaying end point yields spanning two orders of magnitude. Error bars represent standard deviation from three replicate cultures. (C) Sequences of the randomized region (red, panel A) for five T7 promoters used in this study to generate the FrbABCDE library.

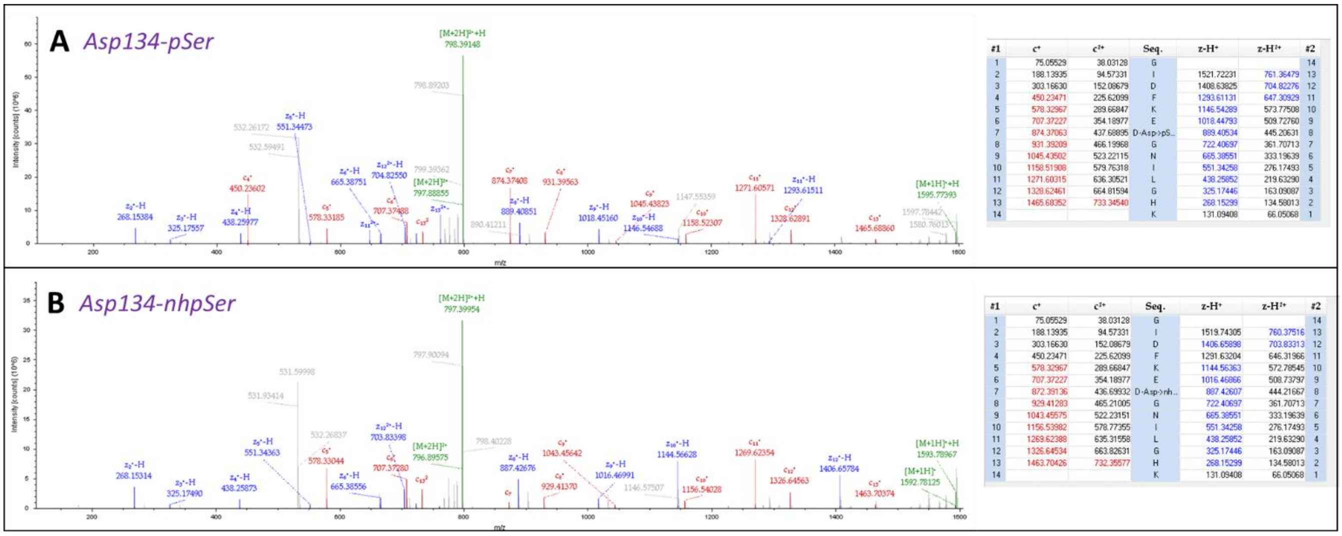


**Supplemental Figure 3**. Bottom-up MS/MS fragmentation of trypsin digested peptides confirming incorporation of (A) pSer at site D134 of 2x-TAG sfGFP made using the pSer GCE system and (B) nhpSer at site D134 of the 2x-TAG sfGFP using the nhpSer Frb-v1 GCE system.

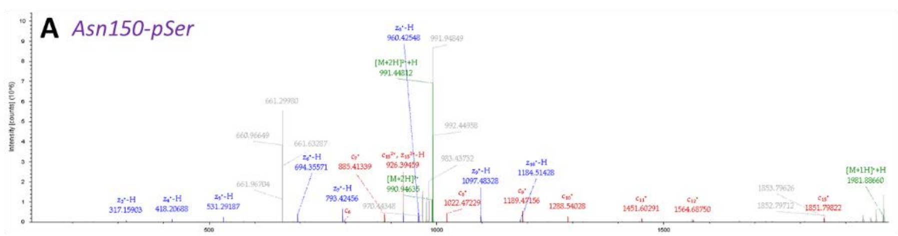


| #1 | c+ | c2+ | Seq. | z-H+ | z-H2+ | #2 |
|---|---|---|---|---|---|---|
| 1 | 131.11789 | 66.06258 | L | | | 16 |
| 2 | 260.16048 | 130.58388 | E | 1851.77111 | 926.38919 | 15 |
| 3 | 423.22381 | 212.11554 | Y | 1722.72852 | 861.86790 | 14 |
| 4 | 537.26674 | 269.13701 | N | 1559.66519 | 780.33623 | 13 |
| 5 | 684.33515 | 342.67121 | F | 1445.62226 | 723.31477 | 12 |
| 6 | 798.37808 | 399.69268 | N | 1298.55385 | 649.78056 | 11 |
| 7 | 885.41011 | 443.20869 | S | 1184.51092 | 592.75910 | 10 |
| 8 | 1022.46902 | 511.73815 | H | 1097.47889 | 549.24309 | 9 |
| 9 | 1189.46738 | 595.23733 | N-Asn->pS... | 960.41998 | 480.71363 | 8 |
| 10 | 1288.53580 | 644.77154 | V | 793.42162 | 397.21445 | 7 |
| 11 | 1451.59912 | 726.30320 | Y | 694.35321 | 347.68024 | 6 |
| 12 | 1564.68319 | 782.84523 | I | 531.28988 | 266.14858 | 5 |
| 13 | 1665.73087 | 833.36907 | T | 418.20582 | 209.60655 | 4 |
| 14 | 1736.76798 | 868.88763 | A | 317.15814 | 159.08271 | 3 |
| 15 | 1851.79492 | 926.40110 | D | 246.12102 | 123.56415 | 2 |

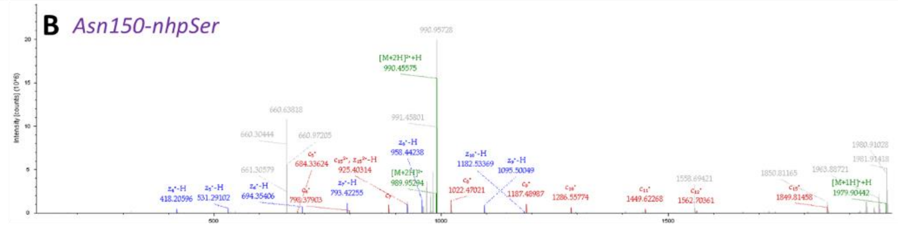


| #1 | c+ | c2+ | Seq. | z-H+ | z-H2+ | #2 |
|---|---|---|---|---|---|---|
| 1 | 131.11789 | 66.06258 | L | | | 16 |
| 2 | 260.16048 | 130.58388 | E | 1849.79185 | 925.39956 | 15 |
| 3 | 423.22381 | 212.11554 | Y | 1720.74926 | 860.87827 | 14 |
| 4 | 537.26674 | 269.13701 | N | 1557.68593 | 779.34660 | 13 |
| 5 | 684.33515 | 342.67121 | F | 1443.64300 | 722.32514 | 12 |
| 6 | 798.37808 | 399.69268 | N | 1296.57459 | 648.79093 | 11 |
| 7 | 885.41011 | 443.20869 | S | 1182.53166 | 591.76947 | 10 |
| 8 | 1022.46902 | 511.73815 | H | 1095.49963 | 548.25345 | 9 |
| 9 | 1187.48812 | 594.24770 | N-Asn->nh... | 958.44072 | 479.72400 | 8 |
| 10 | 1286.55653 | 643.78190 | V | 793.42162 | 397.21445 | 7 |
| 11 | 1449.61986 | 725.31357 | Y | 694.35321 | 347.68024 | 6 |
| 12 | 1562.70392 | 781.85560 | I | 531.28988 | 266.14858 | 5 |
| 13 | 1663.75160 | 832.37944 | T | 418.20582 | 209.60655 | 4 |
| 14 | 1734.78871 | 867.89800 | A | 317.15814 | 159.08271 | 3 |
| 15 | 1849.81566 | 925.41147 | D | 246.12102 | 123.56415 | 2 |

**Supplemental Figure 4**. Bottom-up MS/MS fragmentation of trypsin digested peptides confirming incorporation of (A) pSer at site N150 of 2x-TAG sfGFP made using the pSer GCE system and (B) nhpSer at site N150 of the 2x-TAG sfGFP using the nhpSer Frb-v1 GCE system.

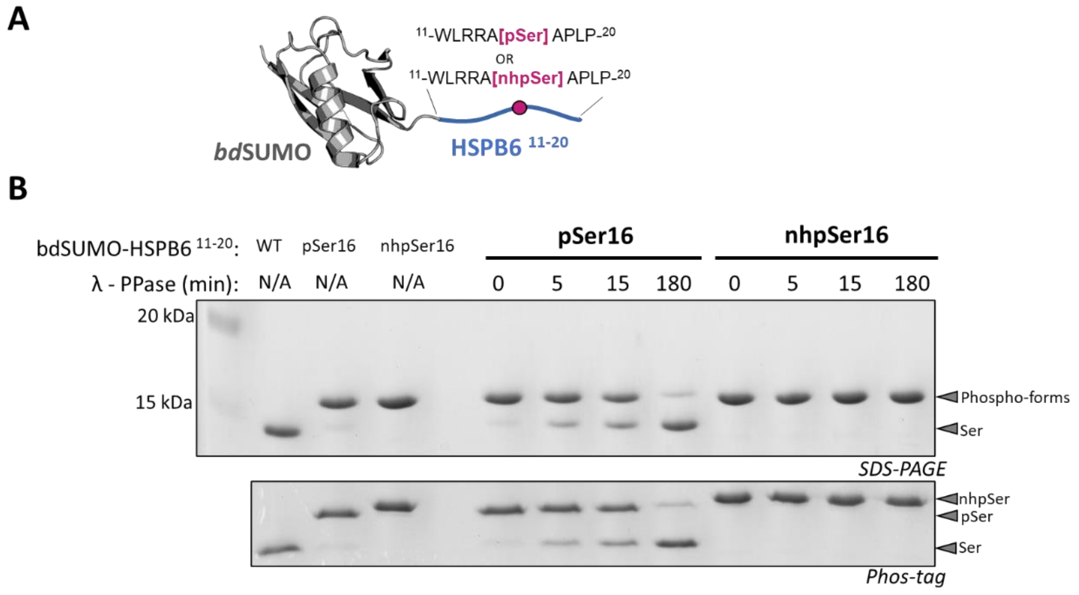


**Supplemental Figure 5**. nhpSer protein made via Frb-v1 GCE is resistant to phosphatase activity. (A) pSer or nhpSer (as well as Ser) was incorporated into site S16 of a HSPB6 peptide consisting of residues 11-20, which was fused at its N-terminus to *bd*SUMO. (B) SDS-PAGE (top) and Phos-tag (bottom) of purified *bd*SUMO-HSPB6$^{11\text{-}20}$ proteins with Ser16, pSer16 and nhpSer16 are shown in the first three lanes. Upon incubation with λ-phosphatase (PPase), the pSer16 containing protein was hydrolyzed as evidenced by it migrating with the electrophoretic mobility of wild-type protein, while the nhpSer protein mobility remained constant. Interestingly, the phospho-proteins migrate more slowly on SDS-PAGE compared to wild-type. Slower electrophoretic mobility of phospho-proteins compared to non-phosphorylated proteins in SDS-PAGE has been well-documented previously.[3] Reasons for this are not well understood but presumably the negatively charged phospho-group limits SDS binding capacity to the protein, lowering its charge to mass ratio and causing it to migrate more slowly.

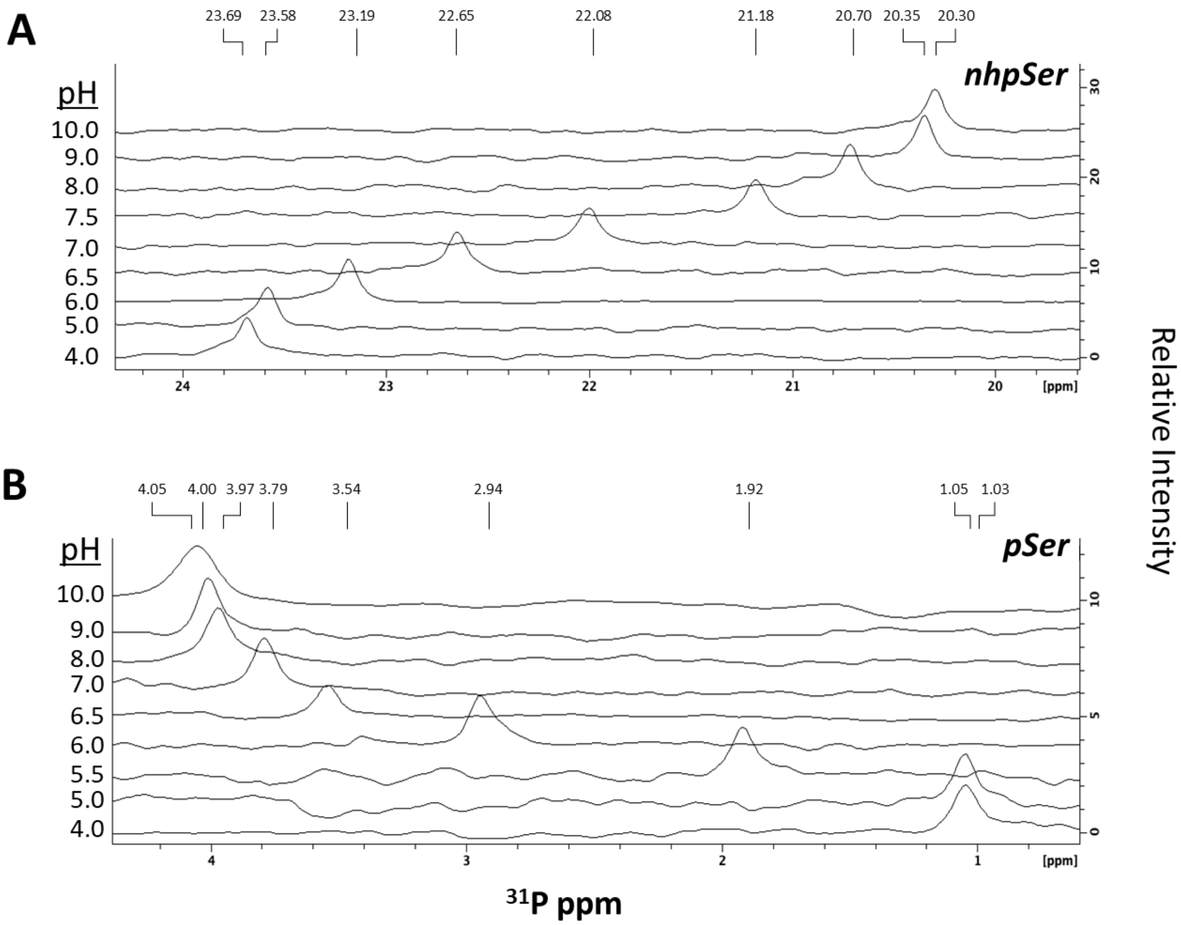


**Supplemental Figure 6**. $^{31}$P NMR spectra of *bd*SUMO-HSPB6$^{11\text{-}20}$ containing (A) nhpSer and (B) pSer at site S16 of HSPB6$^{11\text{-}20}$ at pH's ranging from 4.0 to 10.0. Chemical shifts are indicated above each peak.

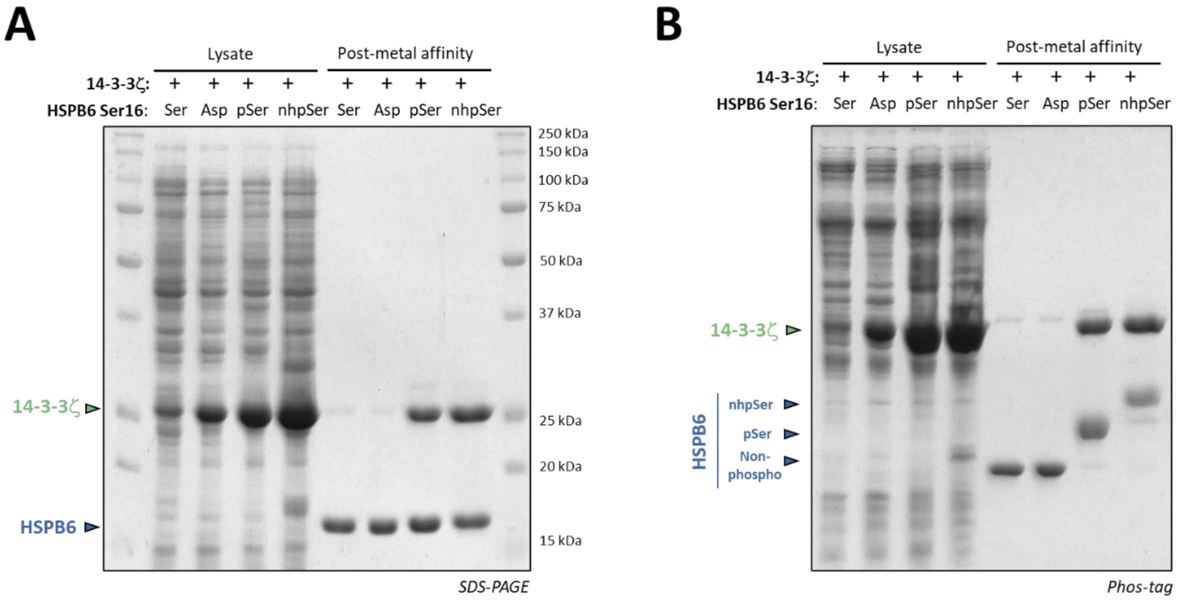


**Supplemental Figure 7**. SDS-PAGE and Phos-tag gel analyses of 14-3-3ζ/HSPB6 (full-length) pull down experiments shown in Fig. 5B. (A) SDS-PAGE of soluble cell lysates confirm expression of 14-3-3ζ in all expressions with HSPB6, indicating that the lack of 14-3-3ζ in the purified wild-type (Ser) and S16D (Asp) HSPB6 samples was due to their inability to complex with 14-3-3ζ. (B) Phos-tag gel of the same samples shown in panel A, confirming pSer and nhpSer incorporation as shown in Fig. 5B.

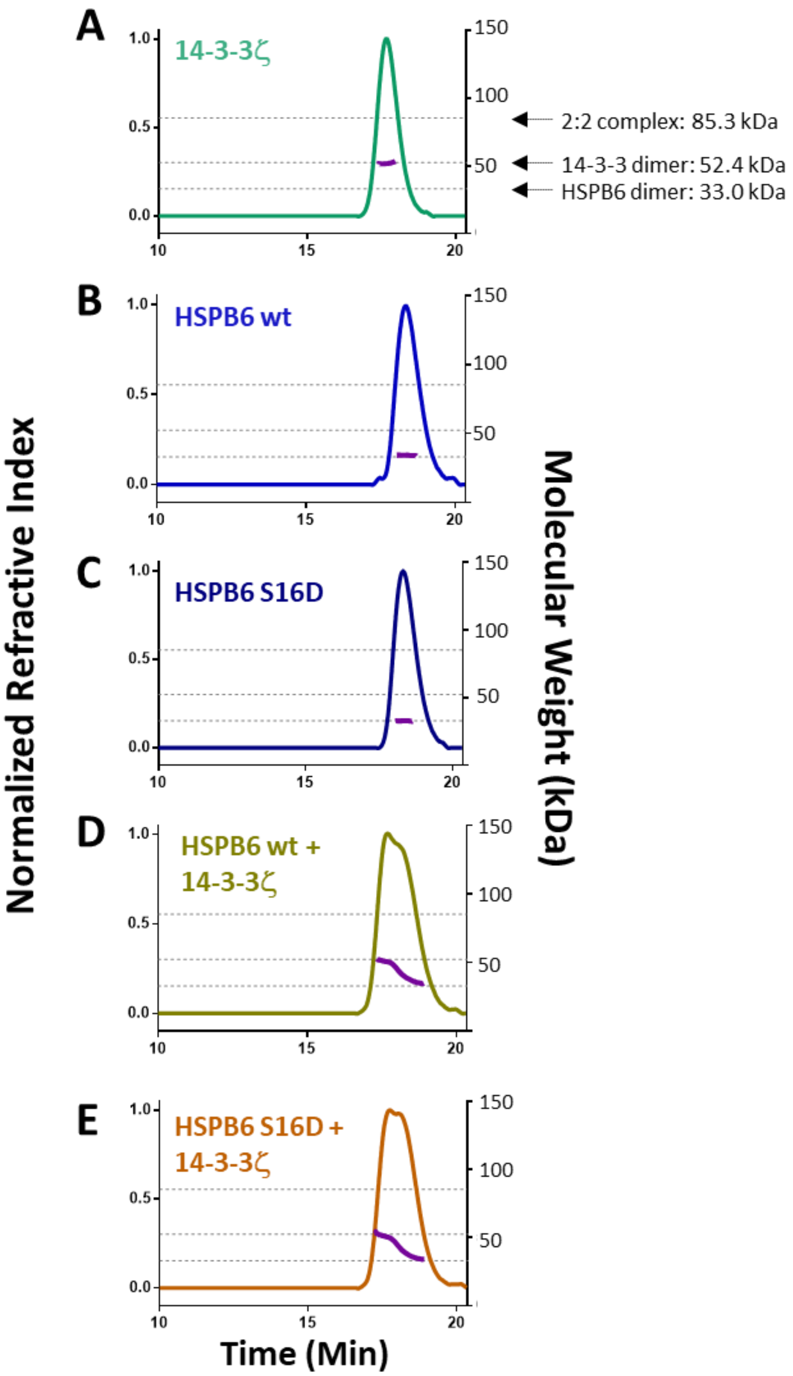


**Supplemental Figure 8**. SEC-MALS analyses of wild-type and S16D HSPB6 with 14-3-3ζ confirm they do not form a stable complex. (A) wild-type 14-3-3ζ only, (B) full-length HSPB6 wt only, (C) HSPB6 S16D, only, (D) an equimolar mixture of 14-3-3ζ and HSPB6 wt, (E) and an equimolar mixture of 14-3-3ζ and HSPB6 S16D. Dotted lines from top to bottom in each panel correspond to the theoretical molecular weights of the 2:2 complex, the 14-3-3ζ dimer and the HSPB6 dimer.

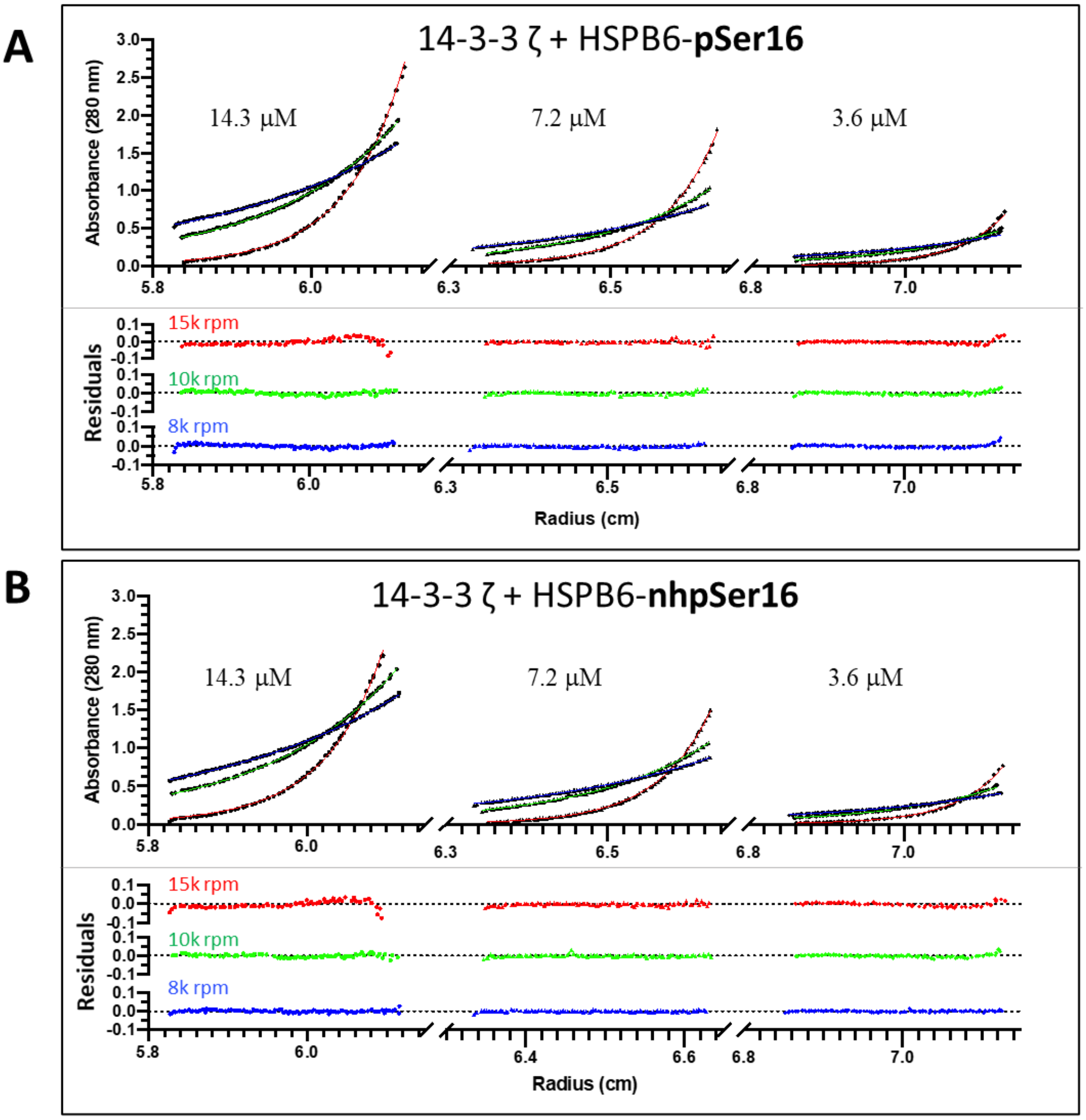


**Supplemental Figure 9**. Equilibrium analytical ultracentrifugation analyses of the (A) pSer16- and (B) nhpSer16-HSPB6/14-3-3 complexes at 14.3 μM concentration (left column), 7.2 μM (middle column) and 3.6 μM (right column) at 15k (red), 10k (green) and 8k (blue) rpm. Residuals from fitting the equilibrium data to a two-state A+B <-->AB model are shown in the bottom row of each panel, in which the pSer and nhpSer complexes fit with dissociation constants of 92 ± 13 and 120 ± 26 nM (errors represent 95% confidence intervals).

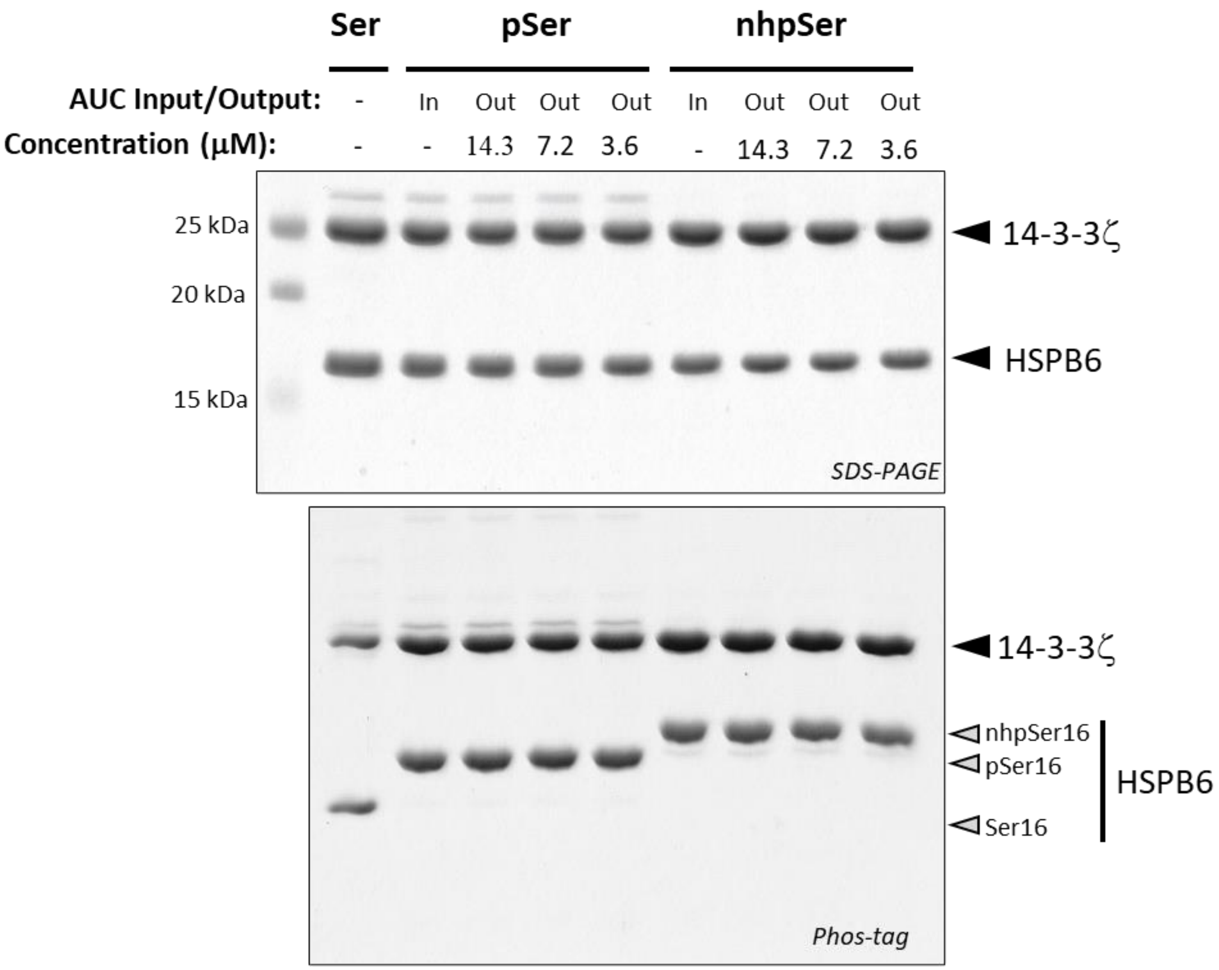


**Supplemental Figure 10.** Evaluation of HSPB6/14-3-3ζ protein sample integrity before and after equilibrium analytical ultracentrifugation (AUC) analysis. Each sample shown in Supplemental Fig. 9 was run after AUC analysis ("out") along side the starting protein sample ("In"). A mixture of purified 14-3-3ζ and wild-type HSPB6 were loaded in the left lane for Phos-tag mobility comparison.

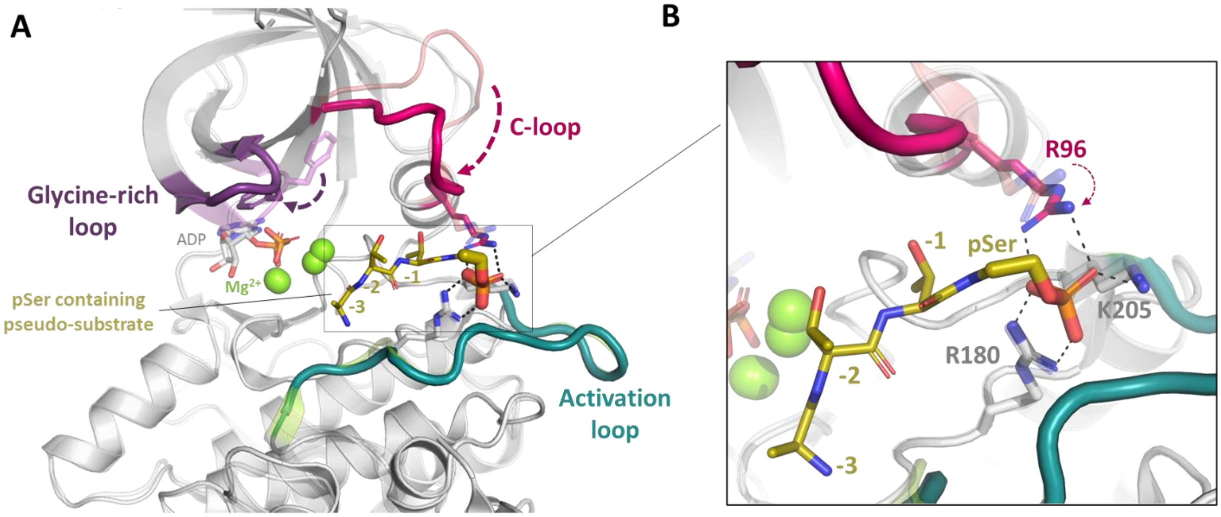


**Supplemental Figure 11**. Mechanism of GSK3β activation upon binding to pSer-containing substrates. (A) Overlayed structures of apo GSK3β (PDB 1pyx) and GSK3β bound to a phosphorylated pseudo-substrate (PDB 4nm3) showing that binding of the peptide (yellow) induces conformational changes in the C-loop (pink) and glycine rich loop (purple). (B) Zoom in showing that the movement of the C-loop upon pSer-substrate binding positions Arg96 to coordinate the phospho group of the substrate peptide, and places the -4 residue (not resolved in crystal structure) in the proximity of the active site where magnesium (green spheres) coordinates ATP.

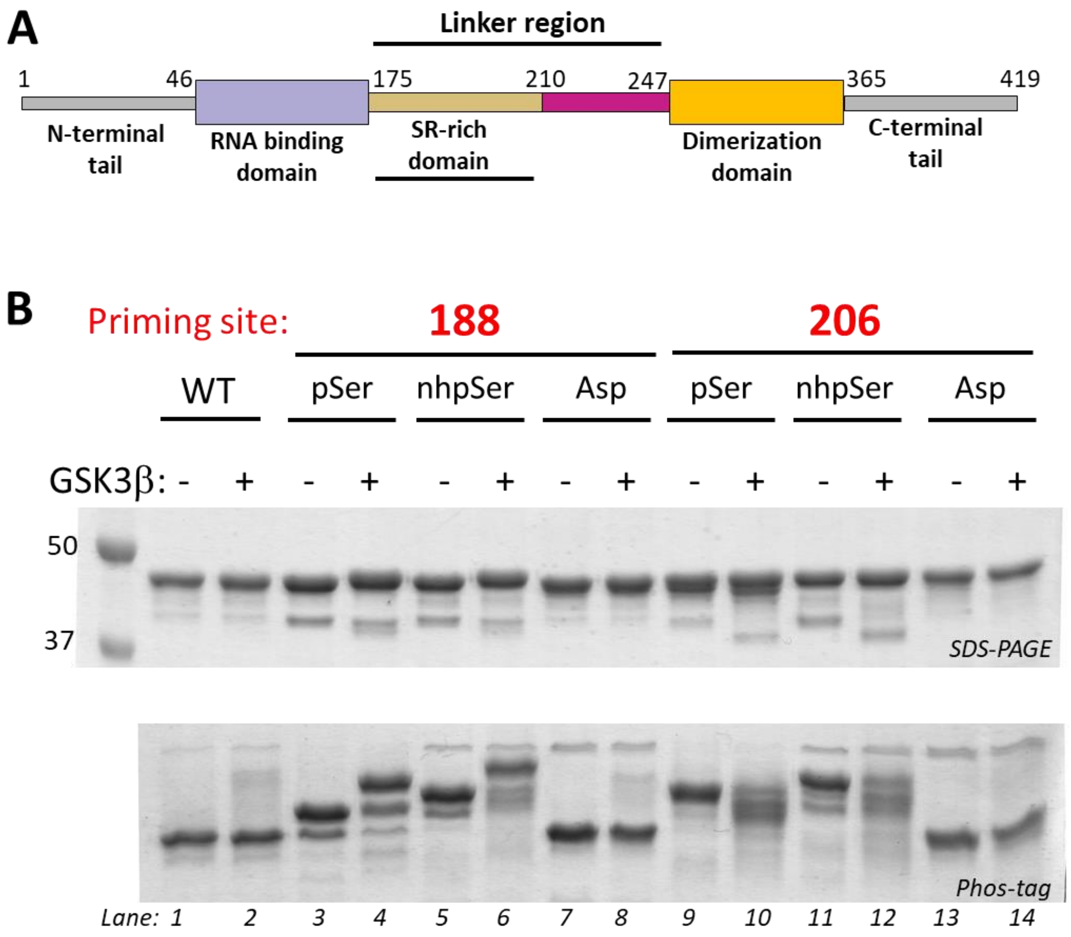


**Supplemental Figure 12**. nhpSer can mimic pSer-dependent GSK3β phosphorylation of full-length Sars-CoV-2 nucleocapsid protein. (A) The SARS-CoV-2 Nucleocapsid Phosphoprotein (Np) contains a Serine-Arginine rich region (SR) in its linker region (Linker-Np, residues 175-247) that connects the N-terminal RNA binding domain and the C-terminal dimerization domain. (B) Ser, Asp, pSer and nhpSer were incorporated at sites S188 and S206 in full-length Np, purified, and mixed with ATP, $Mg^{2+}$, and GSK3β as described in the Methods. SDS-PAGE and Phos-tag gels of reaction products confirm phosphorylation of Linker-Np by GSK3β only in the pSer and nhpSer primed Linker-Np proteins, and not the Ser or Asp proteins, as evidenced by the gel-shifts observed in the Phos-tag gel.

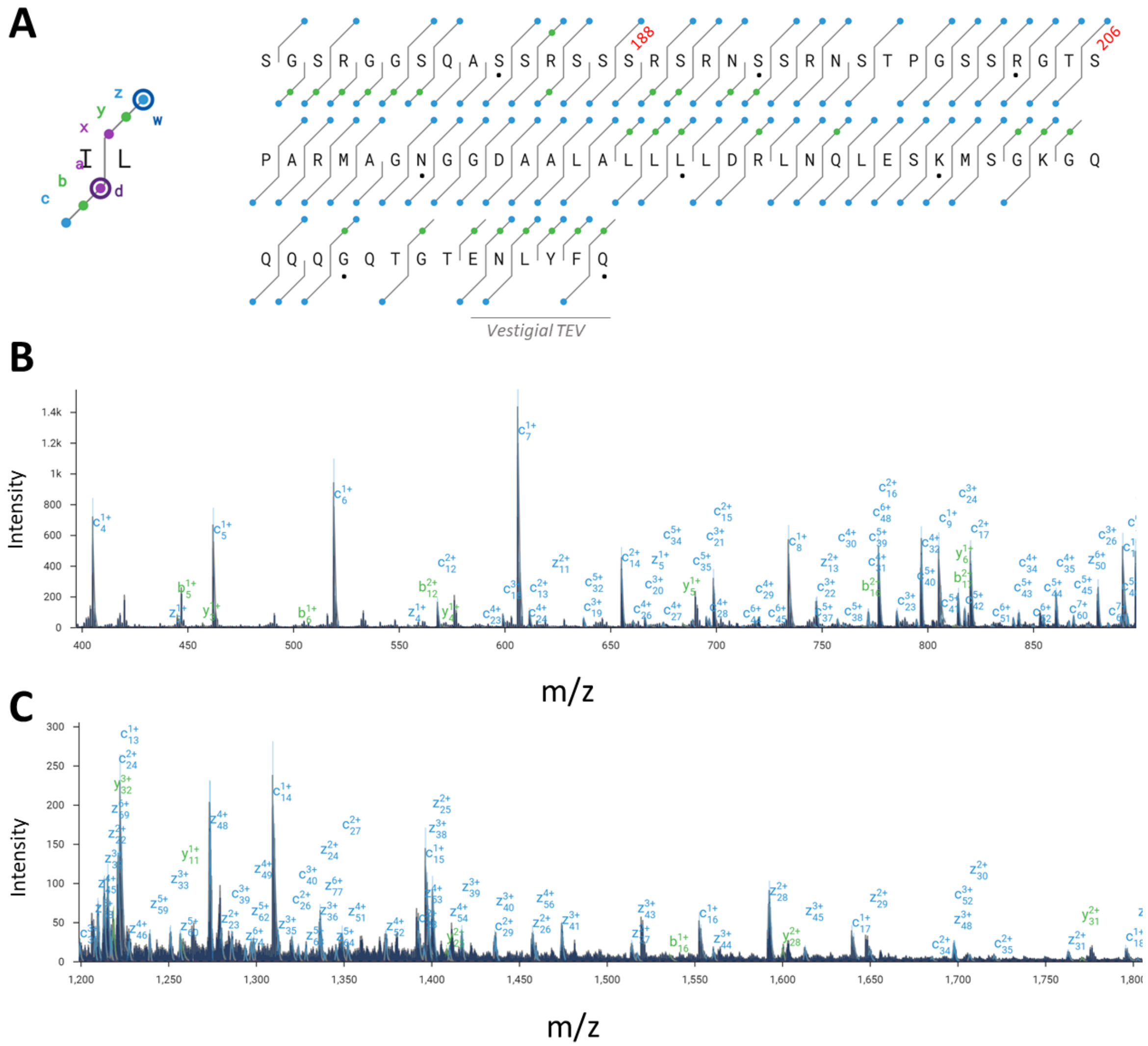


**Supplemental Figure 13**. MS/MS analysis of wild-type Linker-Np protein. (A) Sequence map depicting the detected ion fragments. The blue circles indicate c and z ions, and the green circles indicate b and y ions as shown on the left. Ser188 and Ser206 are labeled accordingly. Block dots represent every 10 residues. The last six residues (ENLYFQ) are part of the TEV protease recognition sequence remaining after removal of sfGFP (Fig. 7C). The top-down analysis yielded an overall sequence coverage of 91%. (B-C). MS/MS fragmentation mass spectra with peaks labeled according to their assigned identity.

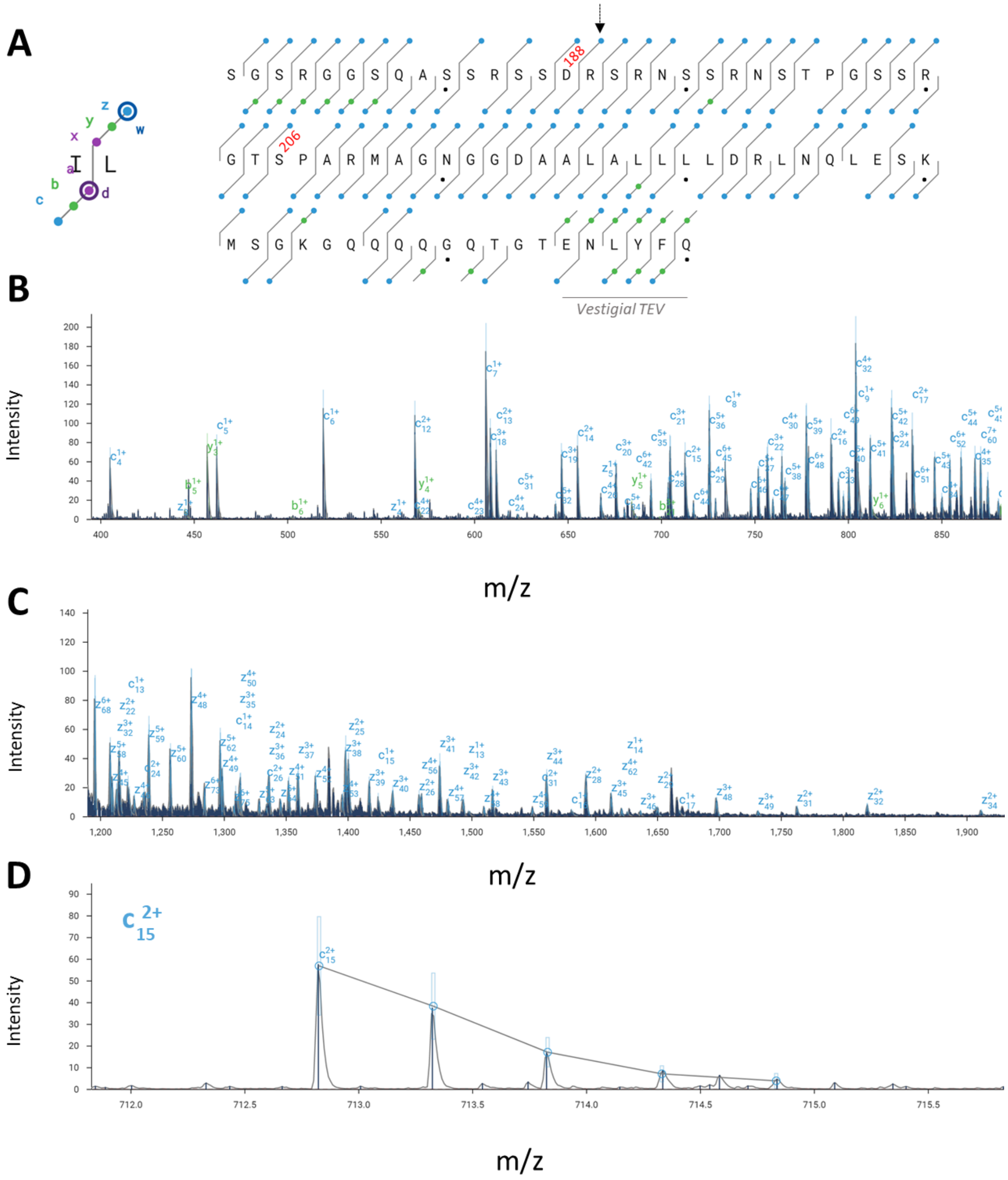


**Supplemental Figure 14**. MS/MS analysis of S188D Linker-Np protein. (A) Sequence map depicting the detected ion fragments. The blue circles indicate c and z ions, and the green circles indicate b and y ions as shown on the left. Asp188 and Ser206 are labeled accordingly. Block dots represent every 10 residues. The last six residues (ENLYFQ) are part of the TEV protease recognition sequence remaining after removal of sfGFP (Fig. 7C). The top-down analysis yielded an overall sequence coverage of 87%. Black arrow points to the fragmented peptide ion shown in panel D. (B-C). Representative MS/MS fragmentation mass spectra with peaks labeled according to their assigned identity. (D) Zoom in of the $c^{2+}/15$ fragmented peptide ion confirms the location of Asp at position 188.

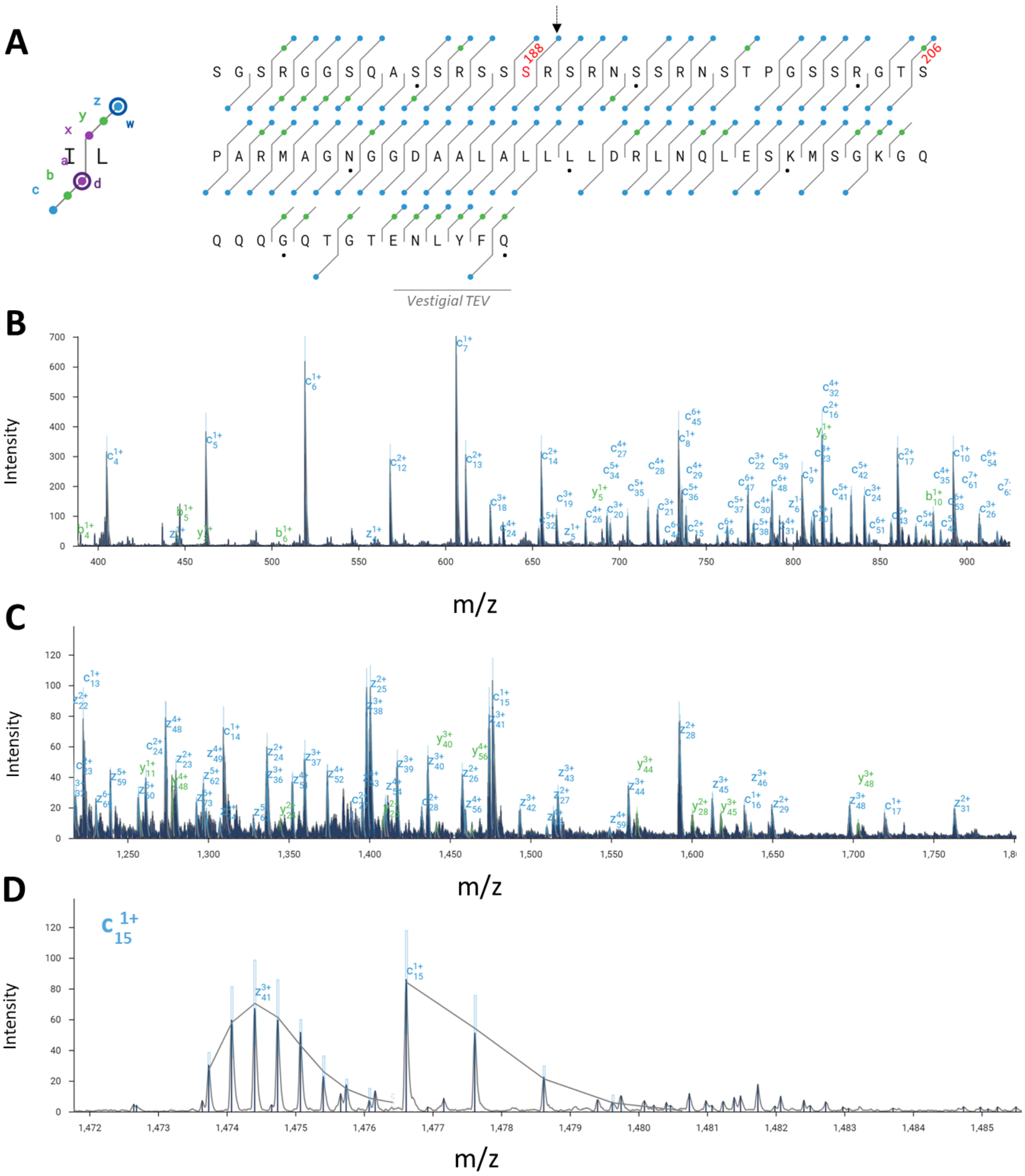

**Supplemental Figure 15**. MS/MS analysis of pSer188 Linker-Np protein. (A) Sequence map depicting the detected ion fragments. The blue circles indicate c and z ions, and the green circles indicate b and y ions as shown on the left. Ser188 and Ser206 positions are labeled accordingly. Block dots represent every 10 residues. The last six residues (ENLYFQ) are part of the TEV protease recognition sequence remaining after removal of sfGFP (Fig. 7C). The top-down analysis yielded an overall sequence coverage of 87%. Black arrow points to the fragmented peptide ion shown in panel D. (B-C). Representative MS/MS fragmentation mass spectra with peaks labeled according to their assigned identity. (D) Zoom in of the $c^{1+}$/15 fragmented peptide ion confirms the location of pSer at position 188.

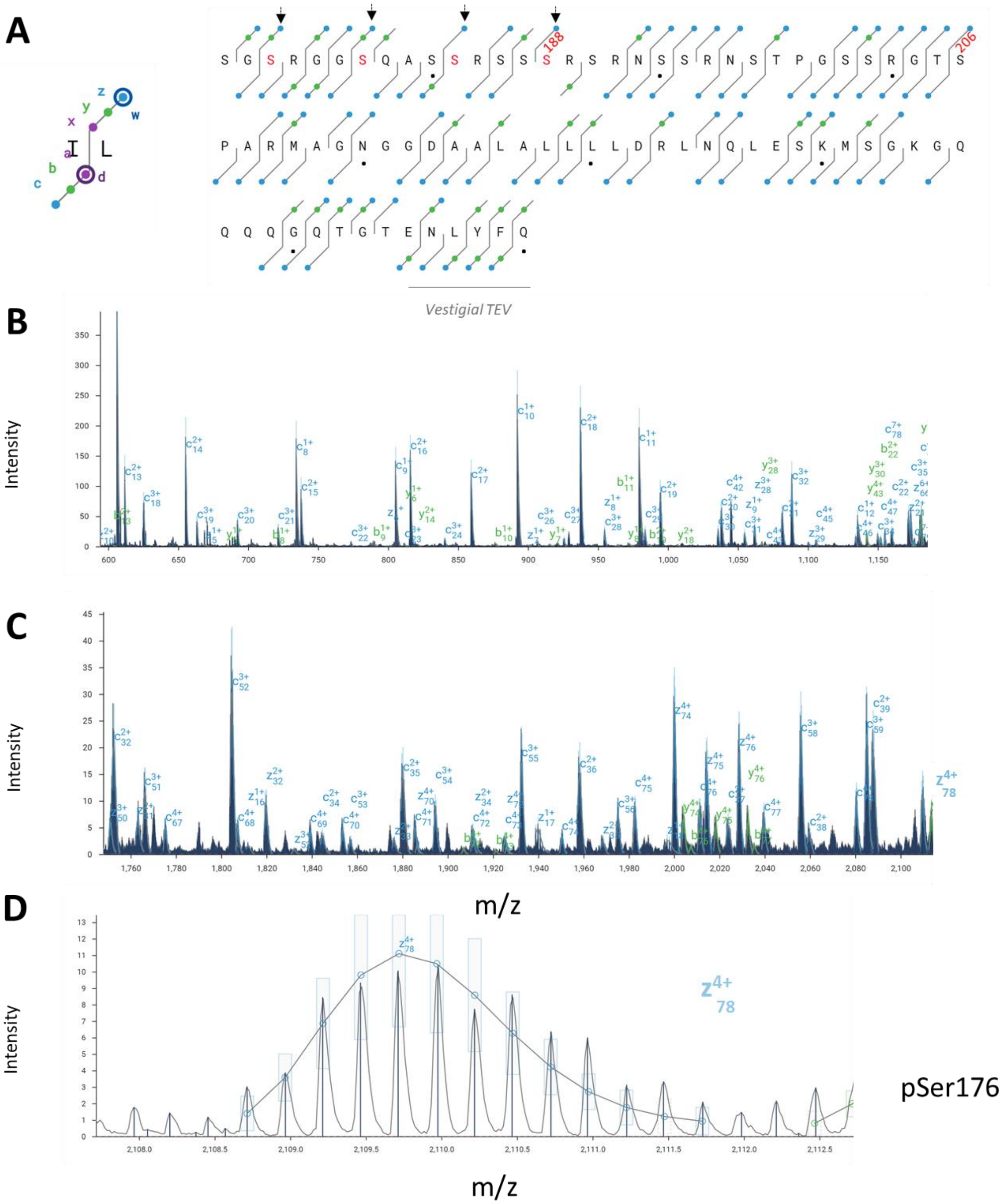


(continued on next page)

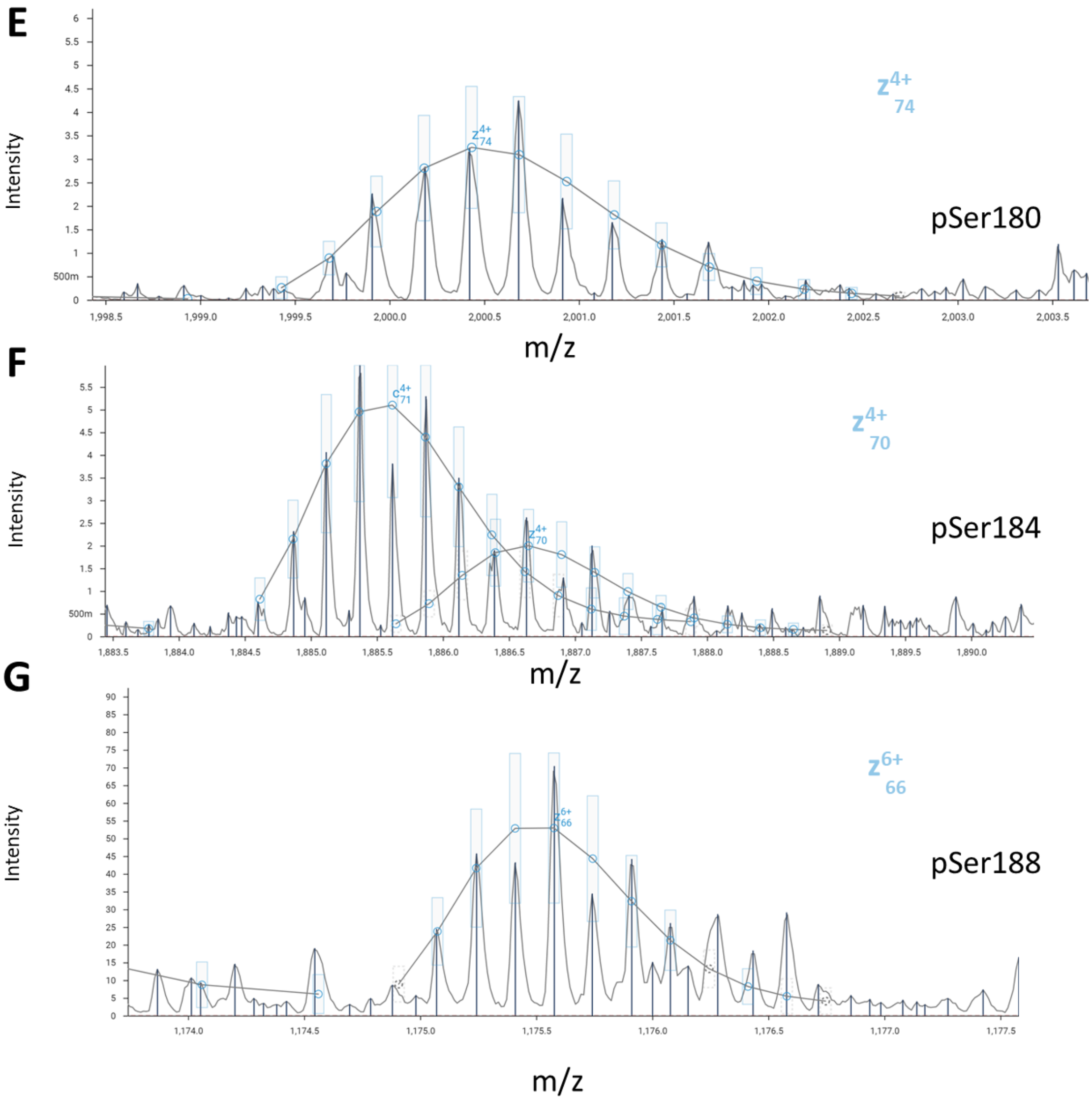


**Supplemental Figure 16**. MS/MS analysis of pSer188 Linker-Np protein after reaction with GSK3β. (A) Sequence map depicting the detected ion fragments. The blue circles indicate c and z ions, and the green circles indicate b and y ions as shown on the left. Ser188 and Ser206 positions are labeled accordingly. Block dots represent every 10 residues. The last six residues (ENLYFQ) are part of the TEV protease recognition sequence remaining after removal of sfGFP (Fig. 7C). The top-down analysis yielded an overall sequence coverage of 80%. Black arrow points to the fragmented peptide ions shown in panels D-G. (B-C). Representative MS/MS fragmentation mass spectra with peaks labeled according to their assigned identity. (D-G) Zoom in of the $z^{4+}$/78, $z^{4+}$/74, $z^{4+}$/70, $z^{6+}$/66 fragmented peptide ion confirms the location of pSer at position 176, 180, 184 and 188, respectively.

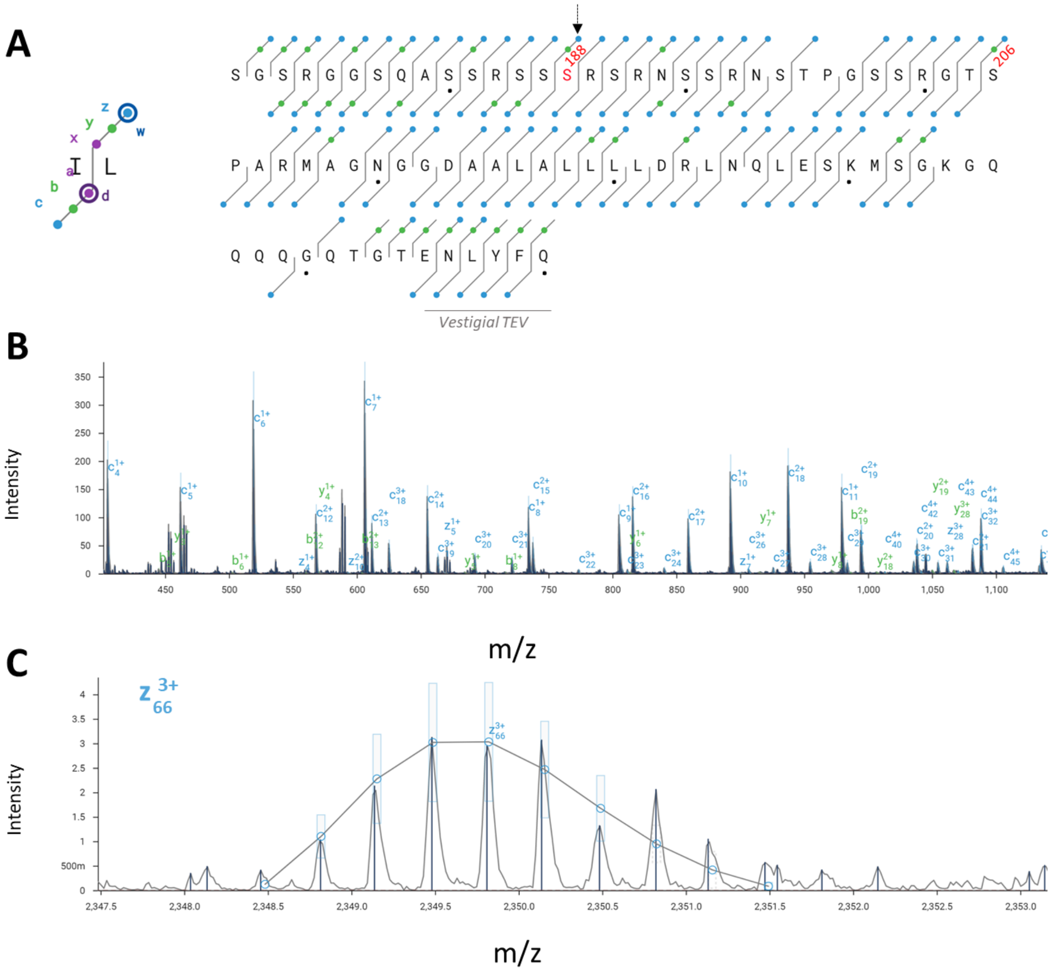


**Supplemental Figure 17**. MS/MS analysis of nhpSer188 Linker-Np protein. (A) Sequence map depicting the detected ion fragments. The blue circles indicate c and z ions, and the green circles indicate b and y ions as shown on the left. Ser188 and Ser206 positions are labeled accordingly. Block dots represent every 10 residues. The last six residues (ENLYFQ) are part of the TEV protease recognition sequence remaining after removal of sfGFP (Fig. 7C). The top-down analysis yielded an overall sequence coverage of 87%. Black arrow points to the fragmented peptide ion shown in panel C. (B). Representative MS/MS fragmentation mass spectra with peaks labeled according to their assigned identity.  (D) Zoom in of the $z^{3+}$/66 fragmented peptide ion confirms the location of nhpSer at position 188.

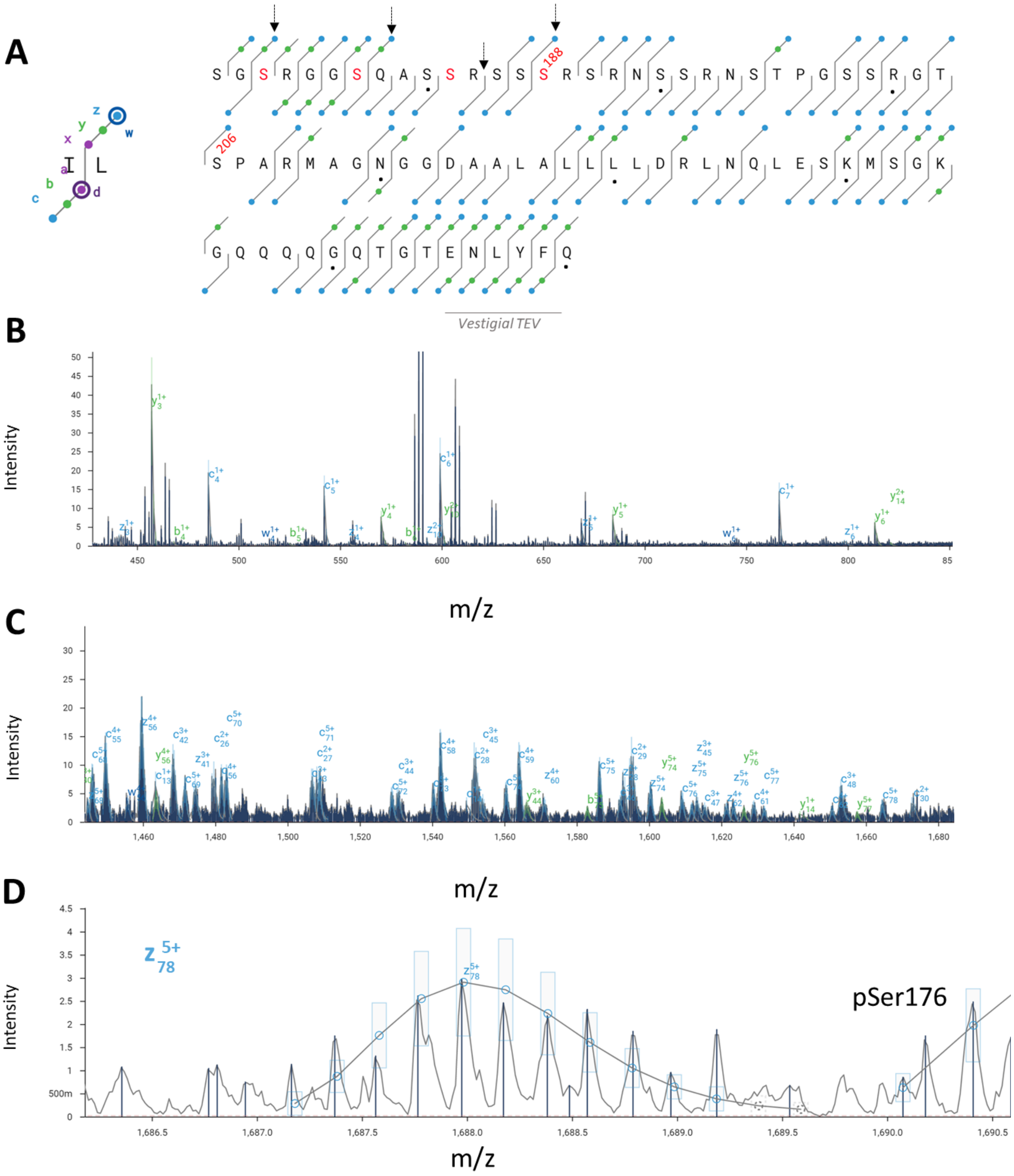


(continued on next page)

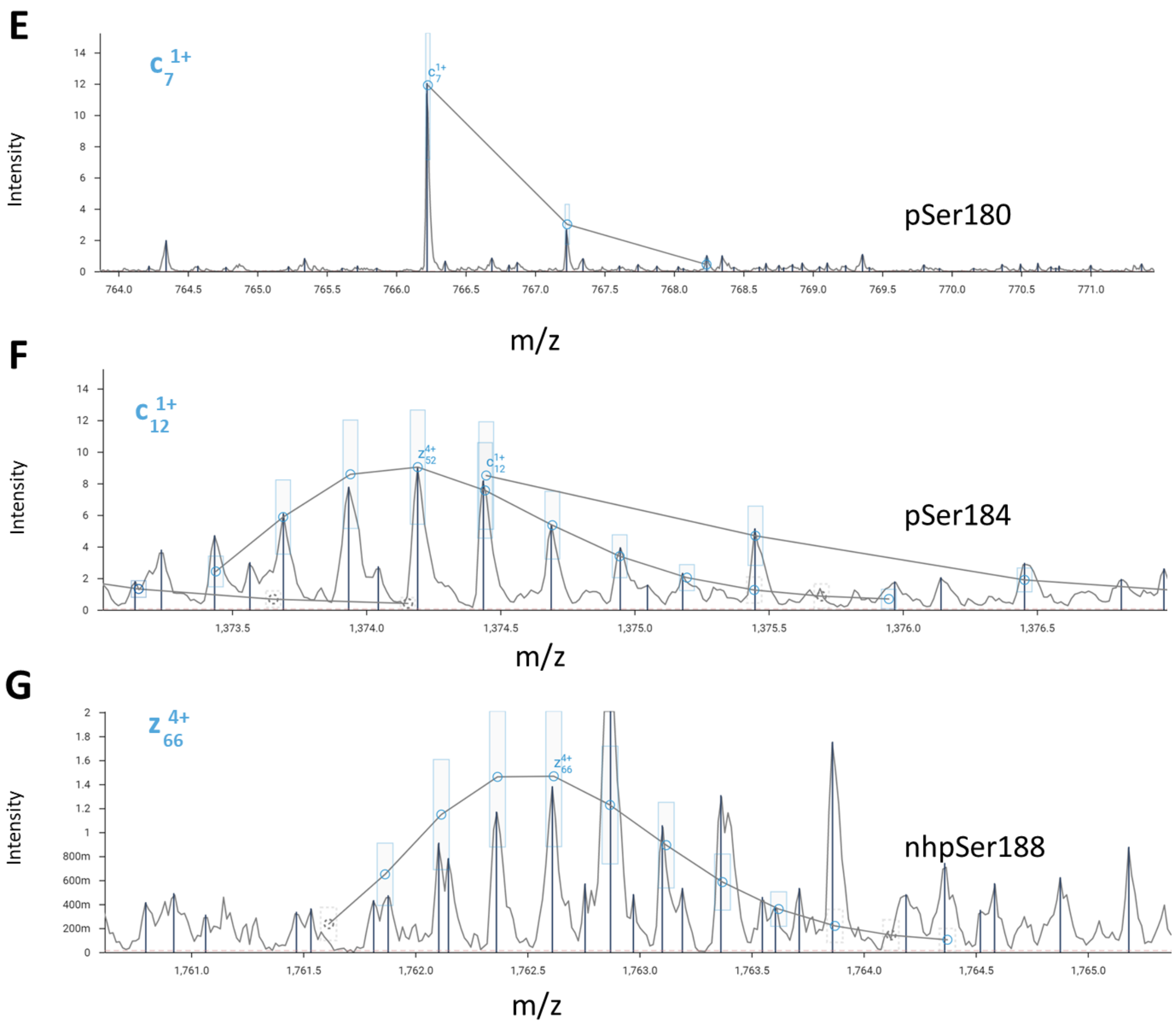


**Supplemental Figure 18**. MS/MS analysis of nhpSer188 Linker-Np protein after reaction with GSK3β. (A) Sequence map depicting the detected ion fragments. The blue circles indicate c and z ions, and the green circles indicate b and y ions as shown on the left. Ser188 and Ser206 positions are labeled accordingly. Block dots represent every 10 residues. The last six residues (ENLYFQ) are part of the TEV protease recognition sequence remaining after removal of sfGFP (Fig. 7C). The top-down analysis yielded an overall sequence coverage of 82%. Black arrow points to the fragmented peptide ions shown in panels D-G. (B-C). Representative MS/MS fragmentation mass spectra with peaks labeled according to their assigned identity. (D-G) Zoom in of the $z^{5+}$/78, $c^{1+}$/7, $c^{1+}$/12, $z^{4+}$/66 fragmented peptide ion confirms the location of pSer at position 176, 180, and 184, and nhpSer at188, respectively.

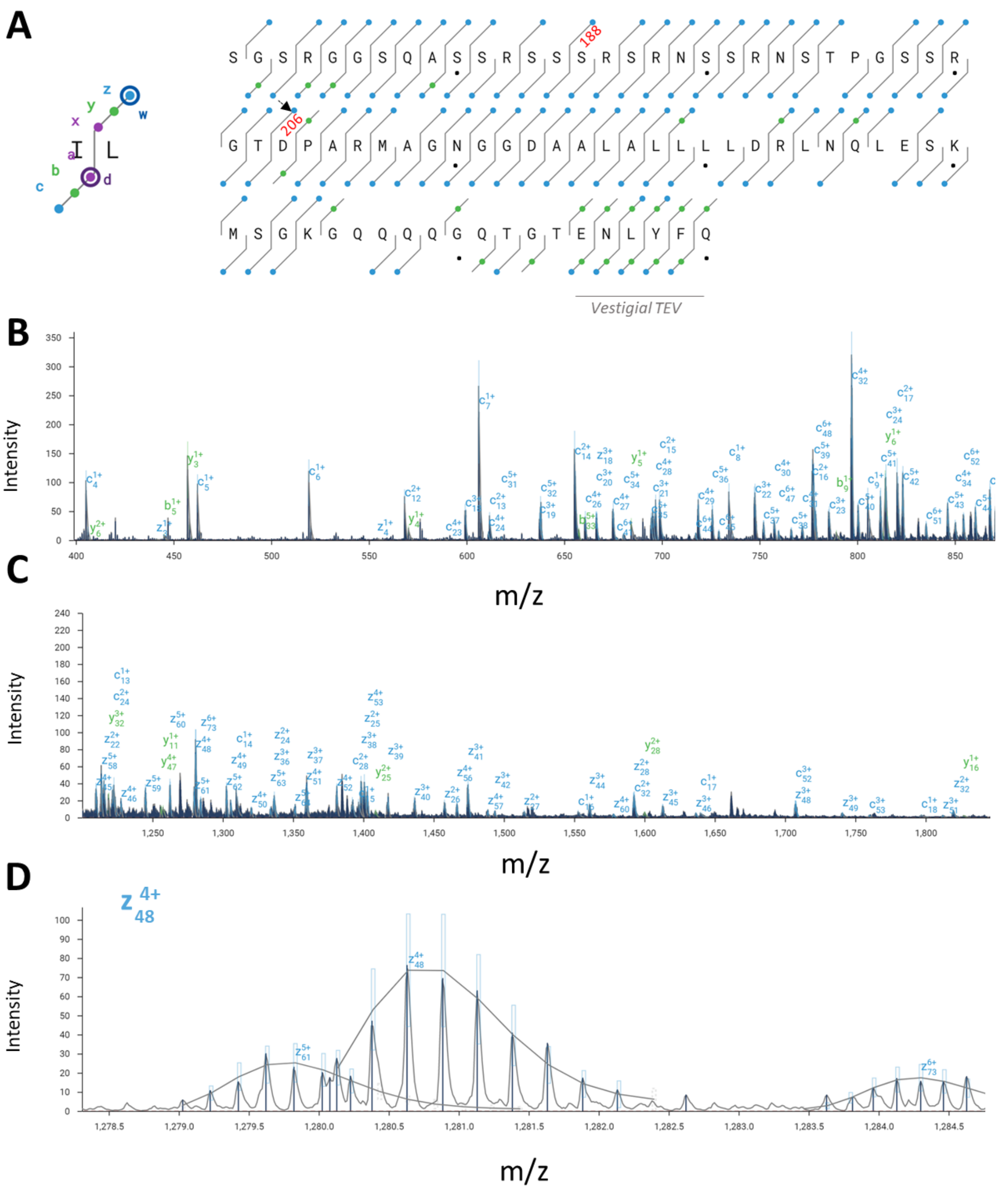


**Supplemental Figure 19**. MS/MS analysis of S206D Linker-Np protein. (A) Sequence map depicting the detected ion fragments. The blue circles indicate c and z ions, and the green circles indicate b and y ions as shown on the left. Ser188 and Asp206 are labeled accordingly. Block dots represent every 10 residues. The last six residues (ENLYFQ) are part of the TEV protease recognition sequence remaining after removal of sfGFP (Fig. 7C). The top-down analysis yielded an overall sequence coverage of 92%. Black arrow points to the fragmented peptide ion shown in panel D. (B-C). Representative MS/MS fragmentation mass spectra with peaks labeled according to their assigned identity. (D) Zoom in of the $z^{4+}$/48 fragmented peptide ion confirms the location of Asp at position 206.

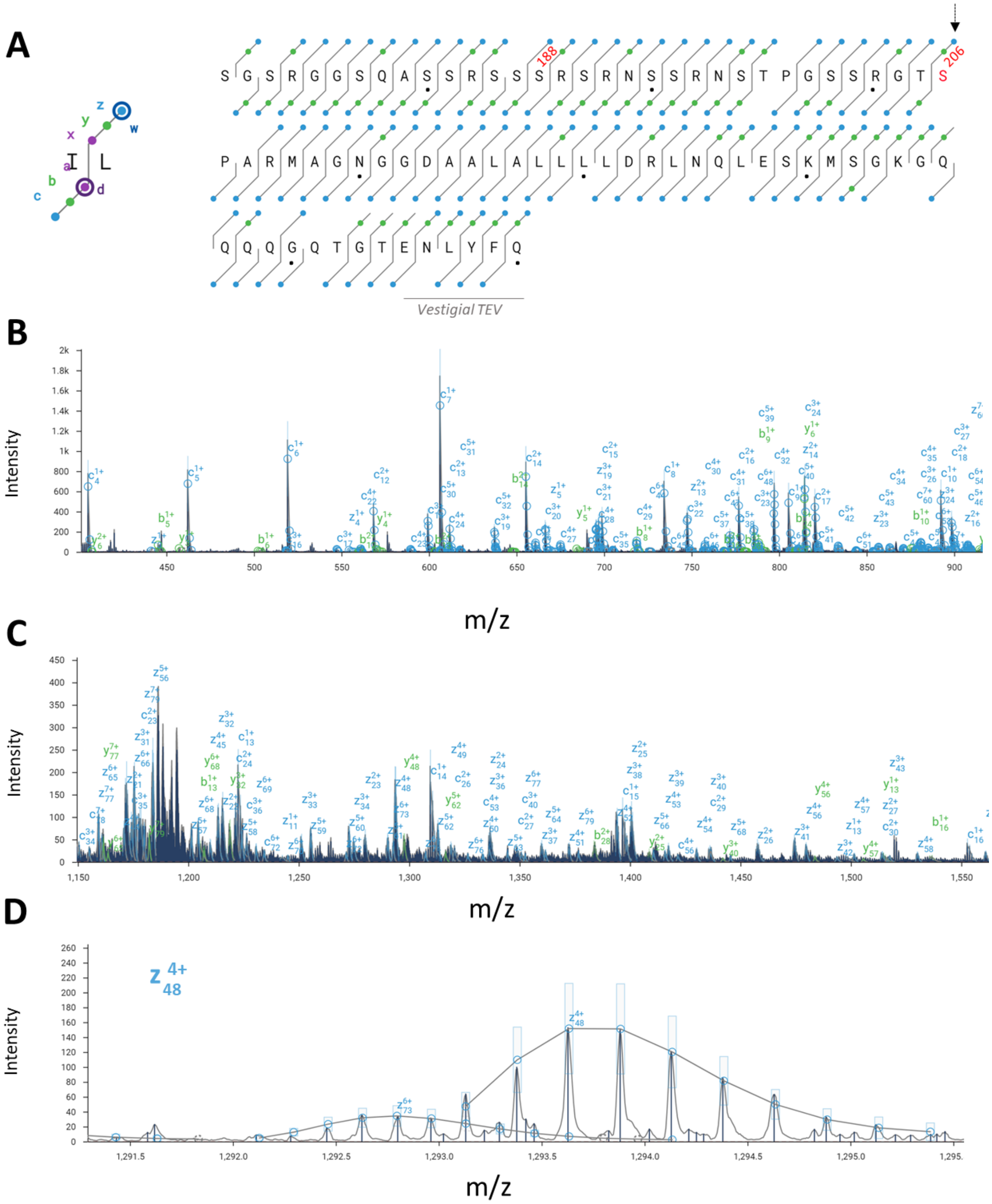


**Supplemental Figure 20**. MS/MS analysis of pSer206 Linker-Np protein. (A) Sequence map depicting the detected ion fragments. The blue circles indicate c and z ions, and the green circles indicate b and y ions as shown on the left. Ser188 and Ser206 positions are labeled accordingly. Block dots represent every 10 residues. The last six residues (ENLYFQ) are part of the TEV protease recognition sequence remaining after removal of sfGFP (Fig. 7C). The top-down analysis yielded an overall sequence coverage of 96%. Black arrow points to the fragmented peptide ion shown in panel D. (B-C). Representative MS/MS fragmentation mass spectra with peaks labeled according to their assigned identity. (D) Zoom in of the $z^{4+}$/48 fragmented peptide ion confirms the location of pSer at position 206.

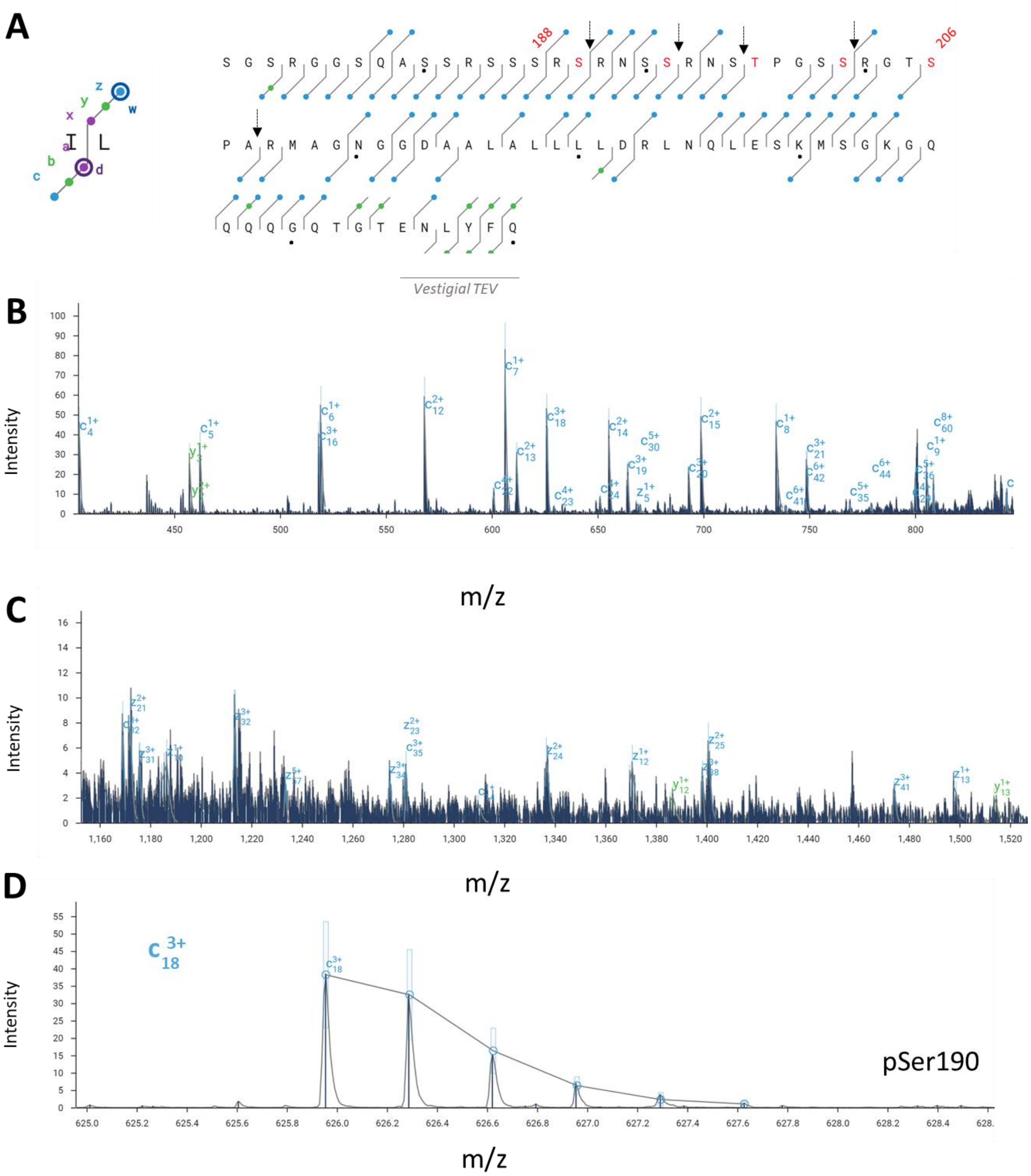


(continued on next page)

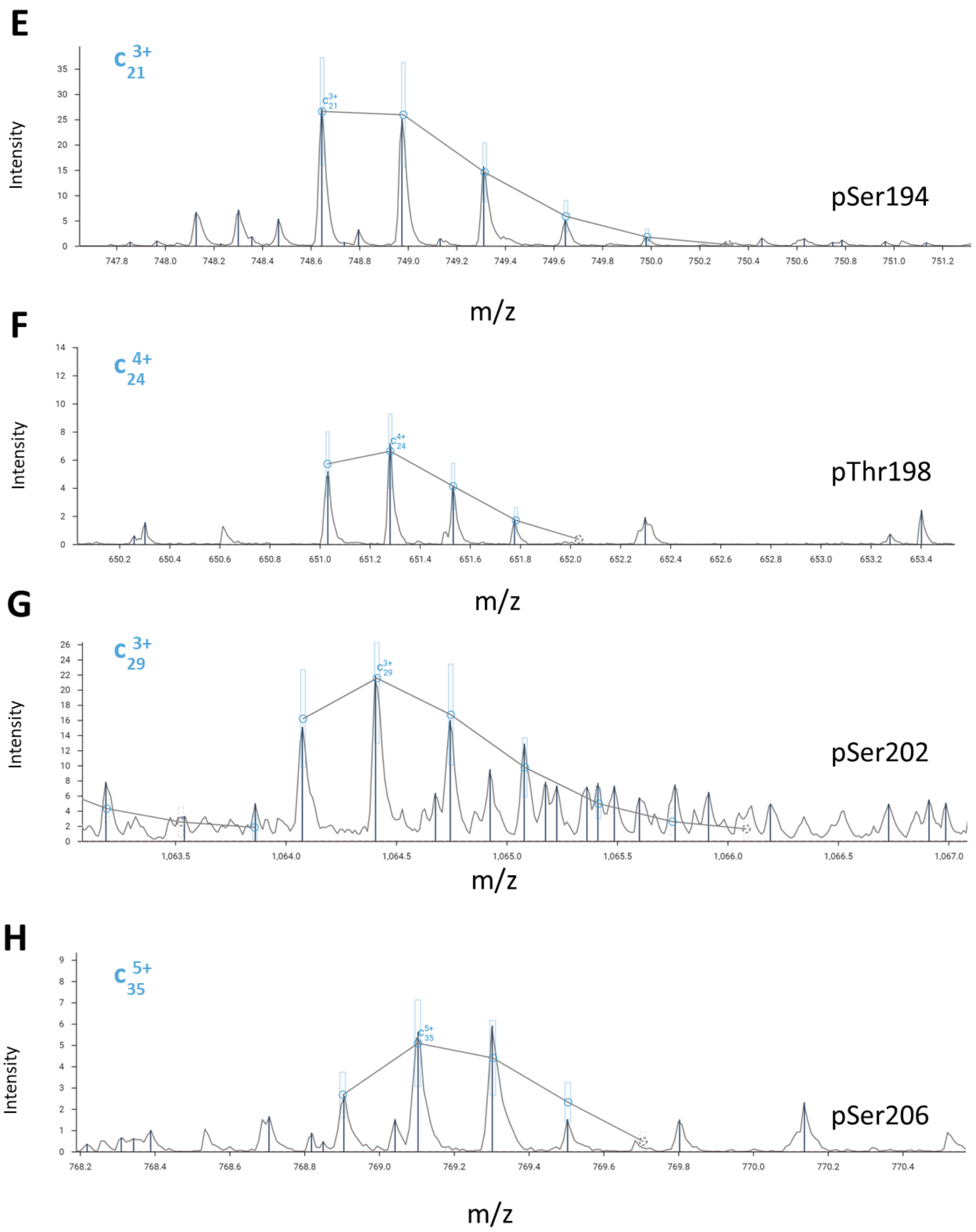


**Supplemental Figure 21**. MS/MS analysis of pSer206 Linker-Np protein after reaction with GSK3β. (A) Sequence map depicting the detected ion fragments. The blue circles indicate c and z ions, and the green circles indicate b and y ions as shown on the left. Ser188 and Ser206 positions are labeled accordingly. Block dots represent every 10 residues. The last six residues (ENLYFQ) are part of the TEV protease recognition sequence remaining after removal of sfGFP (Fig. 7C). The top-down analysis yielded an overall sequence coverage of 78%. Black arrow points to the fragmented peptide ions shown in panels D-H. (B-C). Representative MS/MS fragmentation mass spectra with peaks labeled according to their assigned identity. (D-H) Zoom in of the $c^{3+}$/18, $c^{3+}$/21, $c^{4+}$/24, $c^{3+}$/29, and $c^{5+}$/35 fragmented peptide ion confirms the location of phosphorylation at positions 190, 194, 198, 202 and 206 respectively.

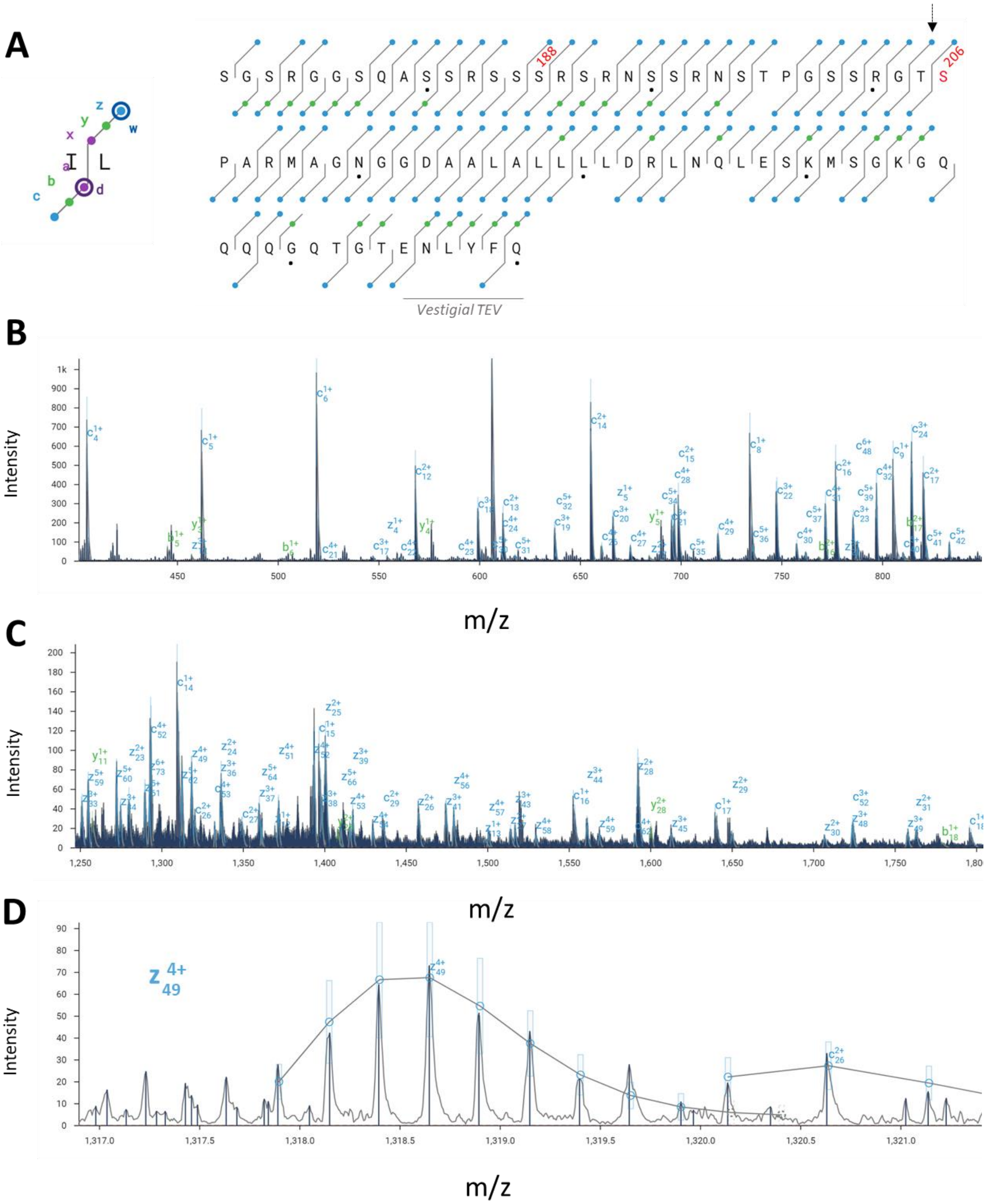


**Supplemental Figure 22**. MS/MS analysis of nhpSer206 Linker-Np protein. (A) Sequence map depicting the detected ion fragments. The blue circles indicate c and z ions, and the green circles indicate b and y ions as shown on the left. Ser188 and Ser206 positions are labeled accordingly. Block dots represent every 10 residues. The last six residues (ENLYFQ) are part of the TEV protease recognition sequence remaining after removal of sfGFP (Fig. 7C). The top-down analysis yielded an overall sequence coverage of 94%. Black arrow points to the fragmented peptide ion shown in panel D. (B-C). Representative MS/MS fragmentation mass spectra with peaks labeled according to their assigned identity. (D) Zoom in of the $z^{4+}$/49 fragmented peptide ion confirms the location of nhpSer at position 206.

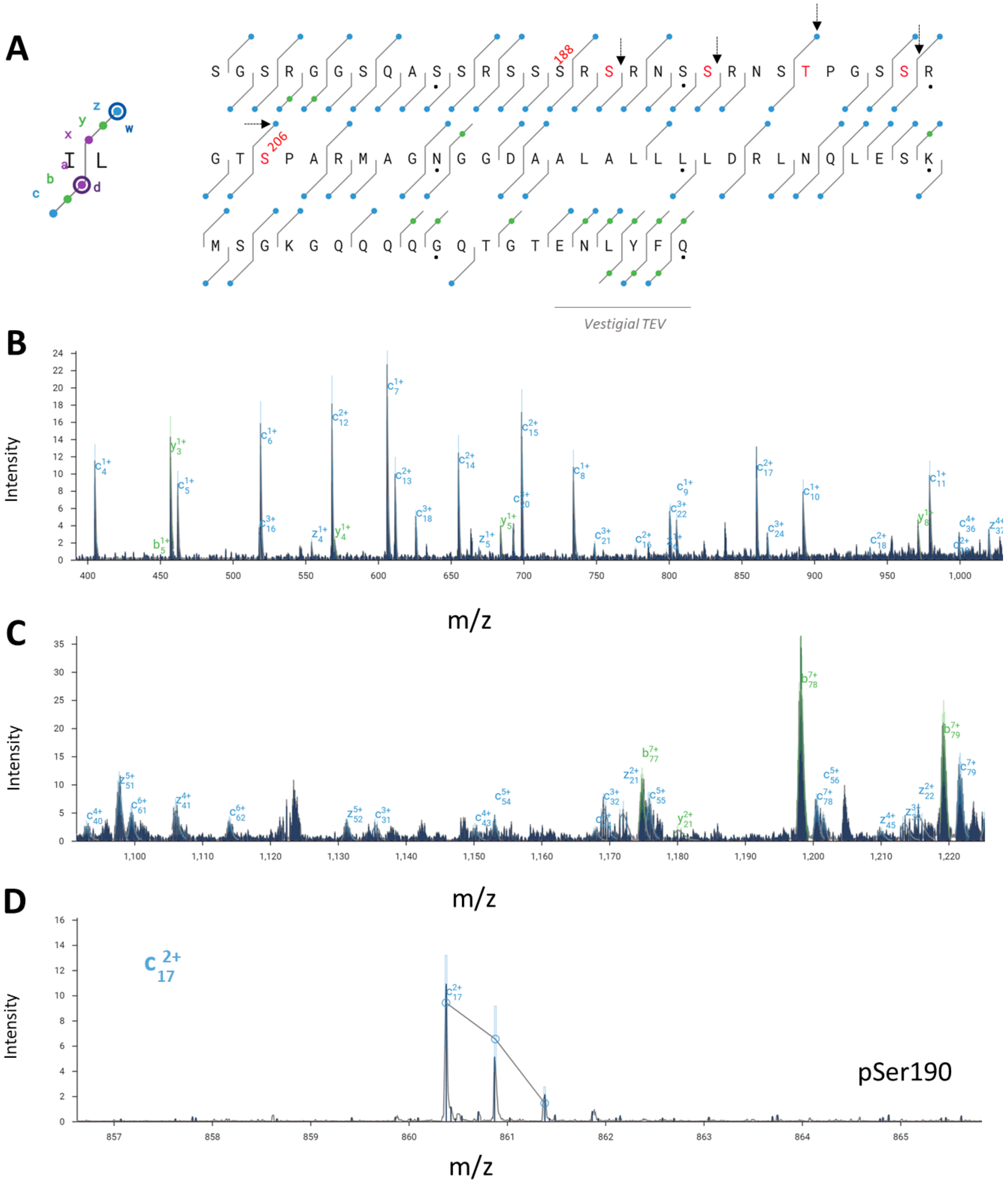


(continued on next page)

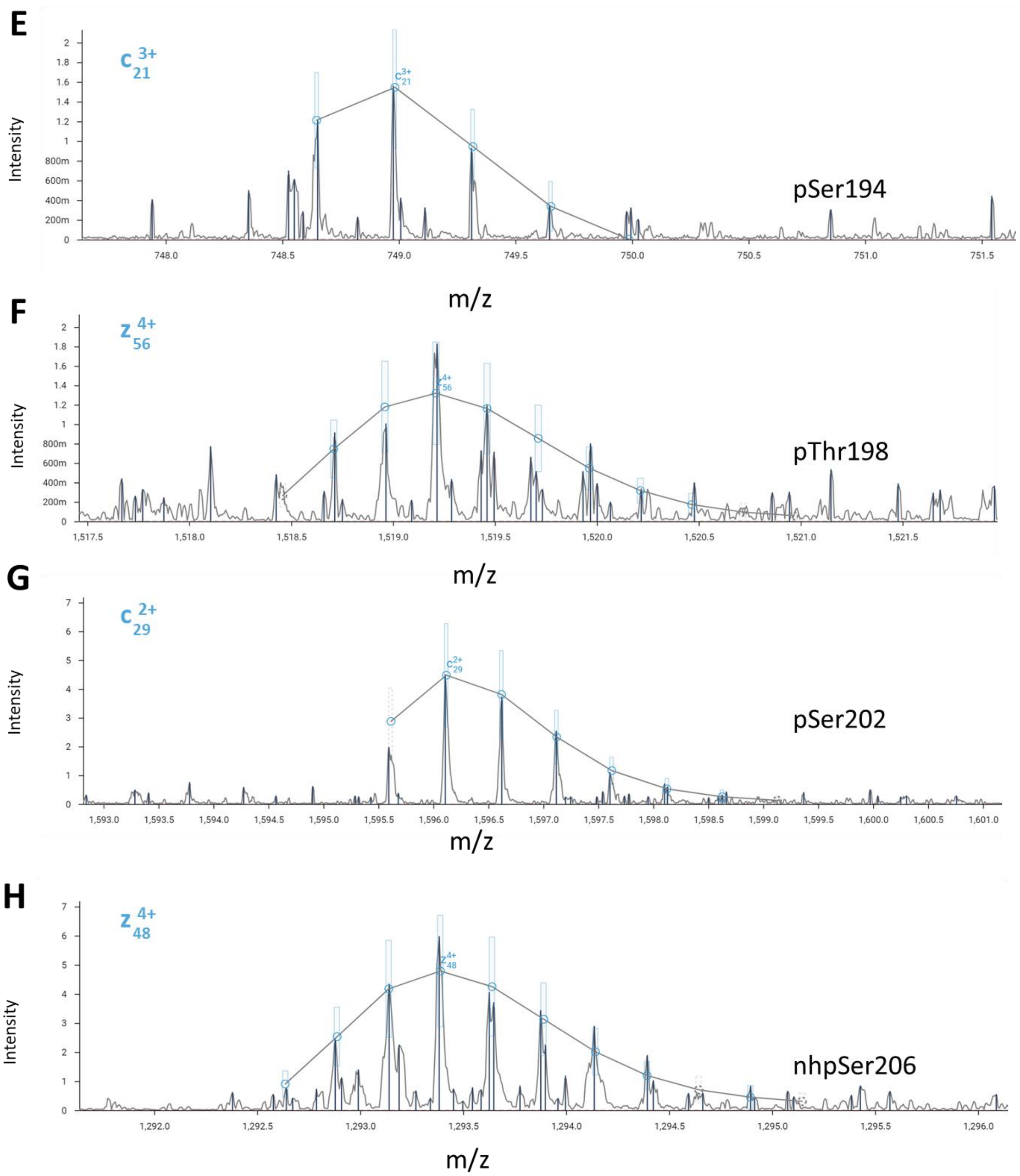


**Supplemental Figure 23**. MS/MS analysis of nhpSer206 Linker-Np protein after reaction with GSK3β. (A) Sequence map depicting the detected ion fragments. The blue circles indicate c and z ions, and the green circles indicate b and y ions as shown on the left. Ser188 and Ser206 positions are labeled accordingly. Block dots represent every 10 residues. The last six residues (ENLYFQ) are part of the TEV protease recognition sequence remaining after removal of sfGFP (Fig. 7C). The top-down analysis yielded an overall sequence coverage of 78%. Black arrow points to the fragmented peptide ions shown in panels D-H. (B-C). Representative MS/MS fragmentation mass spectra with peaks labeled according to their assigned identity. (D-H) Zoom in of the $c^{2+}$/17, $c^{3+}$/21, $z^{4+}$/56, $c^{2+}$/29 and $z^{4+}$/48 fragmented peptide ion confirms the location of phosphorylation at positions 190, 194, 198, and 202, and nhpSer at 206, respectively.

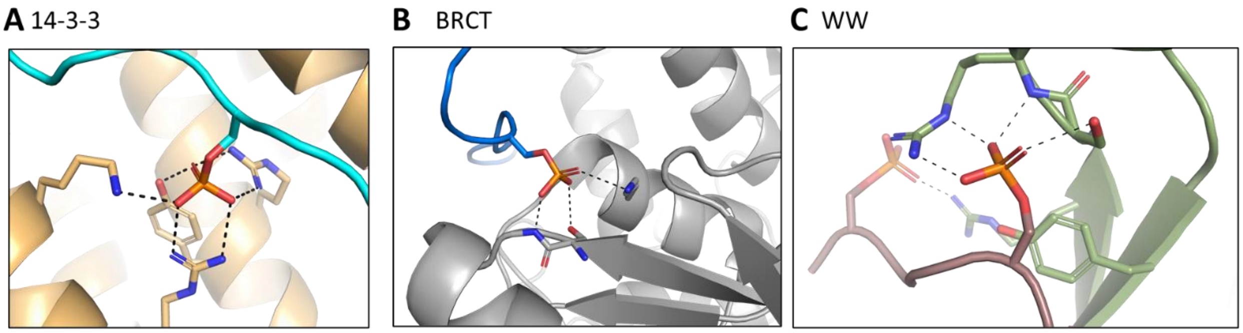


**Supplemental Figure 24**. Prominent phospho-serine binding protein families that do not use the bridging γ-oxygen as a recognition element. Representative structures of (A) 14-3-3 bound to a phosphoserine peptide (PDB 3mhr), (B) a BRCT domain bound to a phosphoserine peptide (PDB 3c0j) and (C) a WW domain bound to a doubly phosphorylated peptide (PDB 1f8a). Each case depicts the structural basis of phosphate-binding specificity for each domain but in each case the pSer bridging γ-oxygen does not make a hydrogen bond with the pSer binding protein. Dotted black lines represent hydrogen bonds.
